# Chronic *Staphylococcus aureus* infection is cured by theory-driven therapy

**DOI:** 10.1101/2020.01.17.910786

**Authors:** Lito A. Papaxenopoulou, Gang Zhao, Sahamoddin Khailaie, Konstantinos Katsoulis-Dimitriou, Ingo Schmitz, Eva Medina, Haralampos Hatzikirou, Michael Meyer-Hermann

**Affiliations:** Department of Systems Immunology and Braunschweig Integrated Centre of Systems Biology, Helmholtz Centre for Infection Research, 38106, Braunschweig, Germany; Institute for Molecular and Clinical Immunology, Medical Faculty, Otto-von-Guericke-University, 39120, Magdeburg, Germany; Systems-oriented Immunology and Inflammation Research Group, Department of Experimental Immunology, Helmholtz Centre for Infection Research, 38124, Braunschweig, Germany; Department of Molecular Immunology, ZKF2, Medical Faculty, Ruhr-University Bochum, 44780, Bochum, Germany; Department of Infection Immunology, Helmholtz Centre for Infection Research, 38124, Braunschweig, Germany; Technische Univesität Dresden, Center for Information Services and High Performance Computing, 01062, Dresden, Germany; Mathematics Department, Khalifa University, P.O. Box: 127788, Abu Dhabi, UAE; Institute for Biochemistry, Biotechnology and Bioinformatics, Technische Universität Braunschweig, 38106, Braunschweig, Germany; Centre for Individualised Infection Medicine (CiiM), 30625, Hannover, Germany

**Keywords:** Chronic *Staphylococcus aureus* infection, mathematical model, myeloid-derived suppressor cells (MDSCs), chronic bacterial infection, model-driven experiments, heat-killed treatment, *S. aureus* infection cured, sterilizing immunity

## Abstract

*Staphylococcus aureus* is considered a dangerous pathogen due to its ability to evade the immune system and resist multiple antibiotics. These evasive strategies lead to difficult-to-treat chronic infections and abscesses in internal organs including kidneys, which are associated with the expansion of myeloid-derived suppressor cells (MDSCs) and their suppressive effect on T cells. Here, we developed a mathematical model of chronic *S. aureus* infection that incorporates the T-cell suppression by MDSCs and suggests therapeutic strategies to eradicate *S. aureus*. We quantified *in silico* a therapeutic protocol with heat-killed *S. aureus* (HKSA), which we tested *in vivo*. Contrary to conventional administration of heat-killed bacteria as vaccination prior to infection, we administered HKSA as treatment, when the hosts were already chronically infected. Our treatment cured all chronically *S. aureus*-infected mice, reduced MDSCs, and reversed T-cell dysfunction by inducing acute inflammation during ongoing, chronic infection without any use of standard treatments that involve antibiotics, MDSC-targeting drugs (chemotherapy), or procedures such as abscess drainage. This study is a proof-of-principle for a treatment protocol against chronic *S. aureus* infection and renal abscesses by repurposing heat-killed treatments, guided and quantified by mathematical modelling. Our mathematical model further explains why previous treatment with inactivated *S. aureus* administered to long-term infected human patients has not led to cure. Overall, our results can have direct relevance to the design of human therapeutics against chronic *S. aureus* infections.

**In brief:** A theory-driven treatment protocol with heat-killed *S. aureus* eradicates *S. aureus*, reduces MDSCs, and reverses T-cell dysfunction *in vivo*.

**Graphical abstract:** 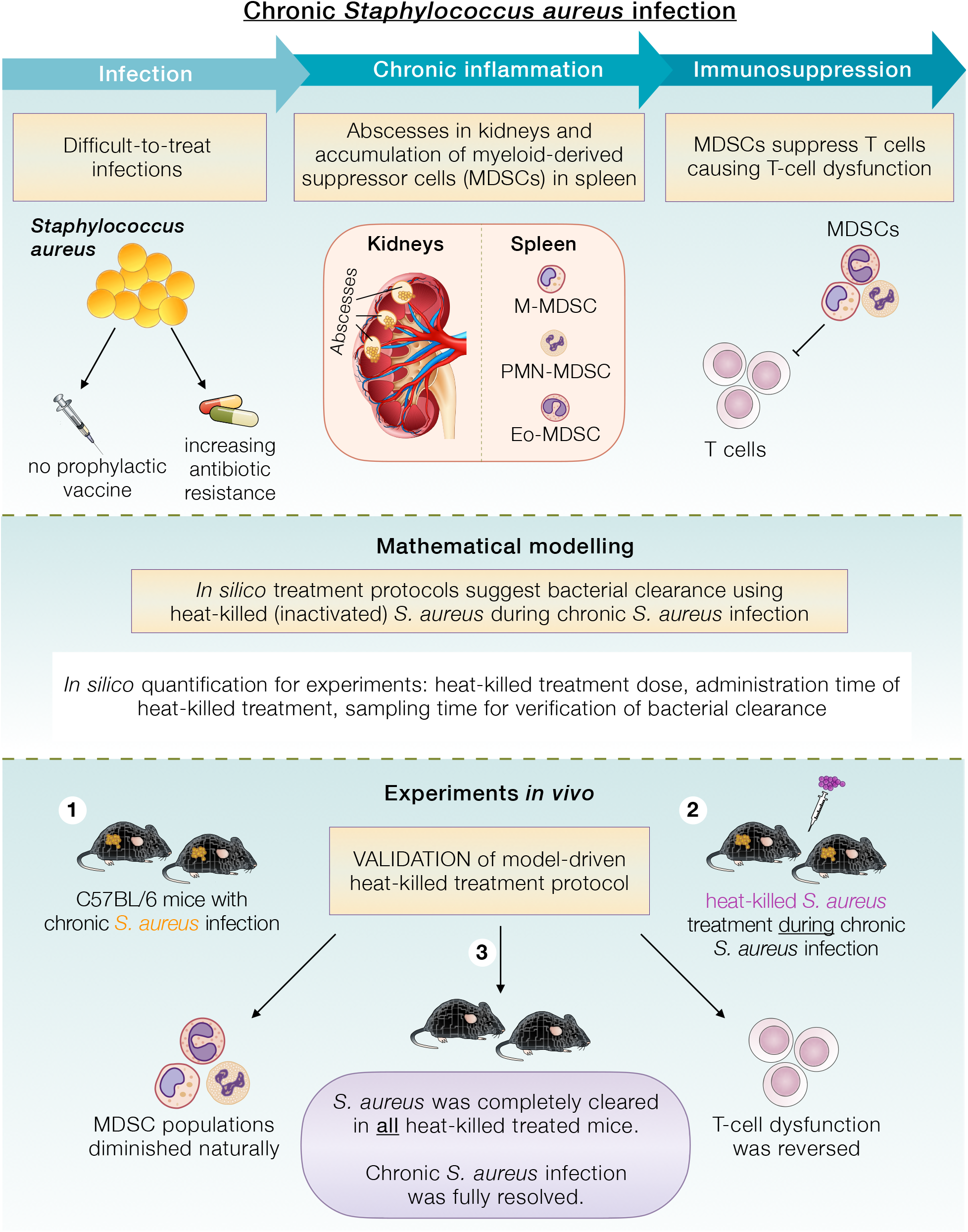

## INTRODUCTION

*Staphylococcus aureus* is a bacterial human pathogen colonizing 20%-30% of the world population and is responsible for nosocomial-acquired and community-acquired infections. Colonization by *S. aureus* as a commensal bacterium is non-problematic. However, after a skin cut, surgery or implantation of medical devices, *S. aureus* can reach deeper tissues and cause life-threatening conditions like pneumonia, endocarditis, osteomyelitis and abscesses in internal organs (*1*, *2*).

The pathogen is of substantial medical concern because it can infect any organ in the body (*1*, *2*), even without being disseminated via the bloodstream (*3*). Moreover, its multiple mechanisms to manipulate and evade immune defenses along with its increasing antibiotic-resistance lead to its persistence in the host and cause chronic, difficult-to-treat infections (*4*). The urgency for new treatments aiming at curing *S. aureus* infections is also emphasized by the WHO, which, on its global priority list of infectious agents, identified *S. aureus* as a “high-priority” pathogen.

Chronic *S. aureus* infections such as chronic osteomyelitis, recurrent furunculosis, and abscesses are hard to eliminate. We have previously shown that during the *chronic phase* of *S. aureus* infection, effectors of innate immunity, such as macrophages (MΦ) and neutrophils, as well as B cells are dispensable for bacterial containment, unlike T cells, which are critical for bacterial control (*5*). However, T cells fail to eradicate the pathogen because prolonged antigenic stimulation (confluent with chronic infection) causes them to enter an anergic state that seems to be irreversible (*5*). As we demonstrated, T-cell dysfunction (also known as T-cell anergy, T-cell suppression, or T-cell hyporesponsiveness) during chronic *S. aureus* infection is attributed to myeloid-derived suppressor cells (MDSCs) rather than other immunosuppressive cells such as regulatory T and B cells, or tolerogenic dendritic cells (*6*).

MDSCs constitute a heterogeneous population of immature myeloid cells that expand in long-lasting pathological conditions, such as chronic bacterial and viral infections (*7*, *8*) including severe SARS-CoV-2 infection (*9*), cancer (*10*), and autoimmunity (*11*). Expansion of MDSCs serves as a natural, anti-inflammatory response to mitigate the detrimental effect of prolonged inflammation (*12*). Often pathogens exploit the immunosuppressive effect by MDSCs to persist within the host, thus establishing chronic infections (*12*). MDSCs are distinguished in three subsets that share the capacity to suppress T cells during chronic *S. aureus* infection: monocytic CD11b^+^Ly6C^+^Ly6G^low^, granulocytic/neutrophilic CD11b^+^Ly6C^low^Ly6G^+^ and eosinophilic CD11b^+^Ly6C^low^Ly6G^low^ MDSCs, known as M-MDSC, PMN-MDSC and Eo-MDSC respectively (*6*, *8*, *13*).

Despite much research, there is no *S. aureus* vaccine to confer prophylaxis against *S. aureus* infections and along with antibiotic-resistance, *S. aureus* is becoming increasingly dangerous every year. It has been estimated that in 2001 *S. aureus* infections afflicted 292045 US hospital inpatients, caused almost 12000 inpatient deaths and cost $9.5 billion in excess charges in US hospitals alone (*14*). By 2014, the number of US inpatients afflicted with methicillin-susceptible and methicillin-resistant *S. aureus* alone has risen dramatically to 616070 individuals and associated costs were estimated to be around $14.6 billion (*15*). Abscesses caused by antibiotic-resistant *S. aureus* have also been increasing. Current treatments rely on simultaneous use of various antibiotics (which promote antibiotic resistance), MDSC-targeting drugs (which are cytotoxic and can have side-effects), and surgery to drain abscesses. Renal abscesses in particular are highly destructive and when multiple antibiotics and abscess drainage fail, immediate nephrectomy is required to save a patient’s life (*16*). Consequently, finding new treatments against *S. aureus* infections is absolutely essential.

Experimental investigations have offered important information on mechanisms of bacterial persistence or MDSC-mediated immunosuppression, however, how to intervene in the complex balance between bacteria, T cells and MDSCs during chronic *S. aureus* infections in order to cure, remains obscure. In this study we constructed a mathematical model to understand the balance between immunity and *S. aureus* during chronic infection and to explore strategies that could cure the infection. Modelling the chronic infection mathematically could bestow a broader observation of possible treatments that would be challenging to discover only by experimental means, whereas *in silico* they could expeditiously and cost-effectively be tested for rendering sterilizing immunity. Mathematical models have been used to shed light on *S. aureus* transmission in community and hospitals (*17*, *18*), staphylococcal growth on foods (*19*), interactions of the pathogen and immune cells during the acute phase of infection (*20*), but also to suggest the optimal sequence of antibiotic administration that could reduce the virulence of *S. aureus* (*21*). However, there are still large gaps on chronic *S. aureus* infections and how to cure them. To our knowledge, this is the first mathematical model that investigates the dynamics of *chronic S. aureus* infection between bacteria and T cells, in the presence of MDSCs, and suggests curative treatments.

Our *in silico* analysis suggested various strategies that could perturb the dynamics of the chronic infection system and cure the infection. For experimental testing *in vivo*, we quantified *in silico* a dose-day treatment protocol using heat-killed (HK), namely inactivated, bacteria. Unlike prior vaccination with HK *S. aureus* (HKSA), which is meant as prophylaxis but instead fails to eradicate the pathogen and exacerbates the infection (*22*), we administered HKSA as treatment, when the hosts were already chronically *S. aureus-*infected. Our *in silico* therapeutic protocol was validated *in vivo*. We here report for the first time reversion of T-cell dysfunction, MDSC-reduction and eradication of *S. aureus* in all HKSA-treated mice without any use of antibiotics, MDSC-targeting drugs, or procedures such as abscess drainage. Our experiments further verified that our HKSA protocol triggered acute inflammation during the already established chronic *S. aureus* infection, which served as the perturbation of the system dynamics and cured the infection. The therapeutic effect of heat-killed treatment is not limited to HKSA, because treatment with heat-killed *Streptococcus pyogenes* (HKSP) also cured a portion of treated animals, reverted T-cell dysfunction and induced acute inflammation during ongoing chronic *S. aureus* infection. Our study is a proof-of-principle for a treatment protocol against chronic *S. aureus* infection and renal abscesses by repurposing heat-killed administration, guided and quantified by mathematical modelling that can have direct relevance to the design of human therapeutics against chronic *S. aureus* infections and abscesses in internal organs.

## RESULTS

### Dynamics of chronic infection

We have previously shown that intravenous inoculation with *S. aureus* results in chronic infection and abscesses in kidneys (*5*, *6*). Bacterial containment in the chronic phase of infection is not attributed to innate immune cells but mainly to effector CD4^+^ T cells, which gradually lose functionality due to suppression by MDSCs (*5*, *6*). Here, we constructed a mathematical model that includes the above-mentioned interactions between bacteria B(t), T cells T(t), and T-cell suppression by MDSCs (Θ) during *chronic S. aureus* infection. In the following the term T cells will refer to CD4^+^ T cells unless otherwise stated.

The ordinary differential equation (ODE) system reads

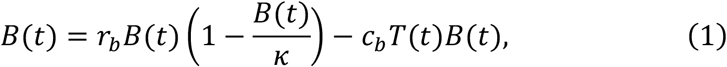

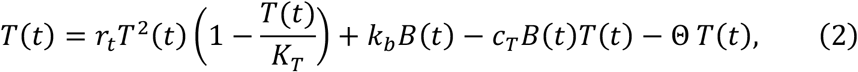

where a dot represents differentiation with respect to time.

During infection, *S. aureus* uses various mechanisms to persist within the host (*23*). This is represented with the term *r_b_B*(*t*)(1−*B*(*t*)/κ) capturing bacterial expansion by logistic growth. Staphylococcal presence stimulates T cells (term *k_b_ B*(*t*)), which proliferate (term *r_t_T*^2^(*t*)). T-cell proliferation is represented with the term *r_t_T*^2^(*t*) because activated T cells secrete Interleukin-2 (IL-2), which induces cell cycle progression of T cells. In return, it creates a positive feedback loop for T-cell proliferation, and hence the term *r_t_T*^2^(*t*) as we reported previously (*24*). The term (1−*T*(*t*)/*KT*) describes the carrying capacity of T cells. As infection becomes chronic, T cells contain bacteria (term *c_b_T*(*t*)*B*(*t*)) (*5*). However, bacterial persistence causes chronic (long-lasting) inflammation, which is harmful to the host. To protect from the adverse effect of prolonged inflammatory signal, MDSCs gradually expand to suppress T-cell activity (*6*). T-cell suppression by MDSCs can happen *systemically,* namely distantly from the site of infection such as in the spleen *(6, 13)*, where T cells increase substantially during chronic *S. aureus* infection (*5*). This is represented with the term Θ *T*(*t*), since MDSC-mediated immunosuppression on T cells requires direct cell–cell contact or cell–cell proximity (*6*). *Locally*, namely at the site of infection, the pathogen can exploit MDSCs towards T-cell suppression to promote its persistence (term *c_T_B*(*t*)*T*(*t*)) *(12)*. A schematic representation of the model is illustrated in Fig. 1A. It is important to note that although innate immune responses do not explicitly appear in the equations, they were not ignored but were rather indirectly incorporated into the term describing bacterial growth, *r_b_B*(*t*)(1−*B*(*t*)/κ) (Eq. (1)). In particular, model parameters were identified with the use of our previously reported experimental results from T and B cells-deficient RAG2^−/−^ mice (only innate immunity present) and immunocompetent mice with chronic *S. aureus* infection and renal abscesses (*5*) (full description in Supplementary Materials). We previously reported major T-cell suppression by MDSCs in spleens of mice with local, chronic *S. aureus* infection in kidneys (*6*, *13*). This phenomenon is known as extramedullar haematopoiesis, happens during chronic inflammation and involves haematopoiesis mostly in spleen, which further induces accumulation of MDSCs in the organ (*25*). In accordance, our fitting revealed that parameter *c_T_* describing local T-cell suppression by MDSCs in kidneys (the site of infection), was negligible compared to parameter Θ describing systemic T-cell suppression by MDSCs, such as in spleens (Supplementary Materials).

**Fig. 1.**
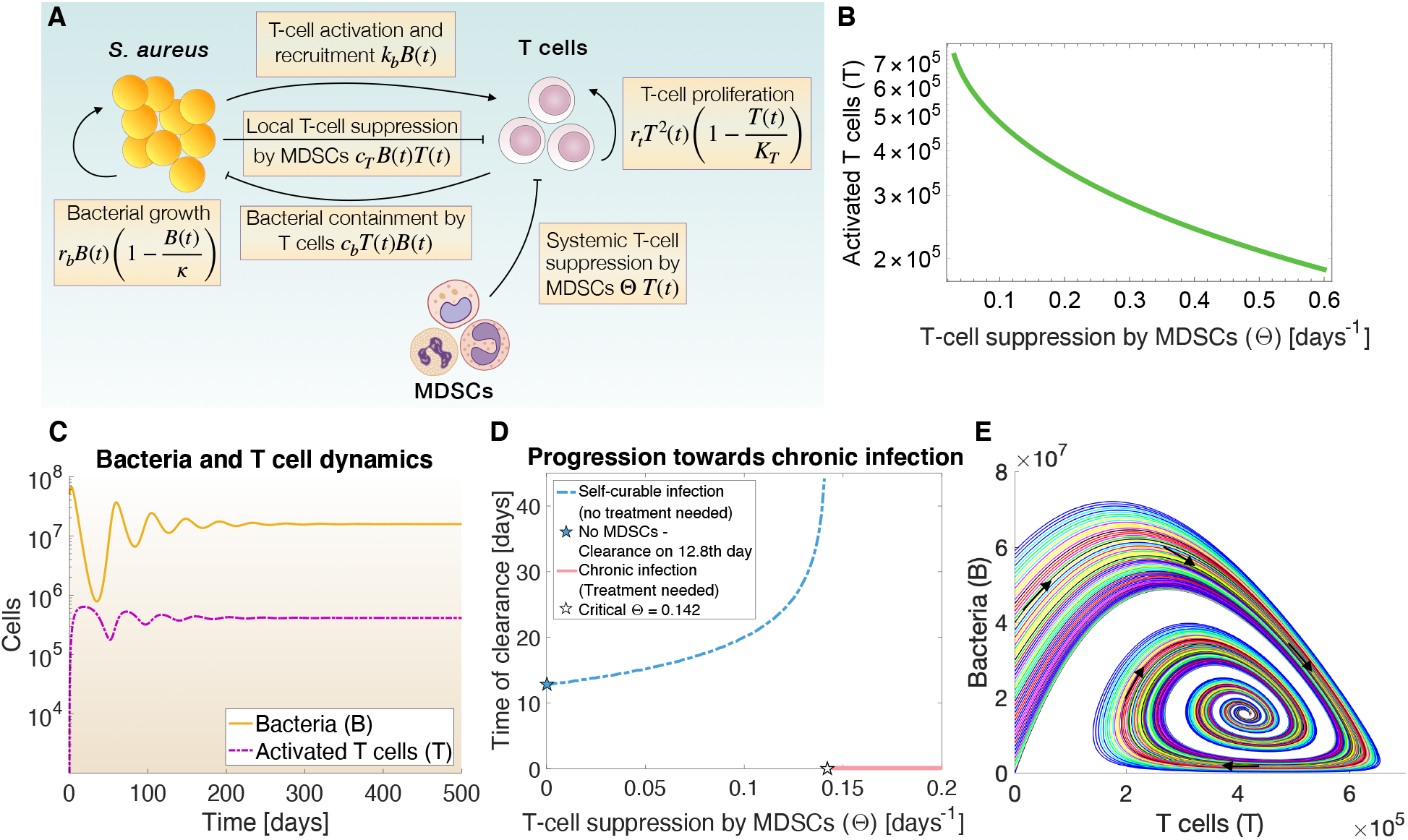
Dynamics of chronic *Staphylococcus aureus* infection. (**A**) Schematic representation of chronic *S. aureus* infection model. (**B**) Correlation between T cells and T-cell suppression by MDSCs (Θ) is inversely proportional. The correlation was plotted using the analytical solutions of the T-cell differential equation in steady state (Supplementary Materials, Eq. (S5)) for increasing values of parameter Θ. (**C**) Dynamics of bacteria (B) and activated T cells (T) in time are shown as numerical solutions of the ODE system. Infection was induced by setting the bacterial population equal to 5×10^7^ cells at day 0 in Eq. (1) to be consistent with experimental bacterial inoculation causing chronic *S. aureus* infection (*6*). Model parameters are as shown in table S1. (**D**) Infection was initiated as in (**C**). Numerical calculation of bacteria for changing values of parameter Θ (T-cell suppression by MDSCs) in the range [0, 0.2]. For bacterial numbers <0.000001, the infection is considered cured (blue), else persisting (pink) and the corresponding day of bacterial clearance is shown or set to zero, respectively. The blue star represents a scenario of MDSC-absence and hence non-existent T-cell suppression (Θ=0). As Θ gradually increases, the infection progresses towards chronic phase and bacterial clearance becomes more difficult. The blue line represents when, despite MDSC-mediated suppression, the infection can be cured by the immune response alone. The white star represents the critical value of Θ, when T cells become anergic by MDSC-mediated suppression, the infection persists and external treatment-intervention is required for cure. table S1 shows the values for the rest of the model parameters. (**E**) Stable steady state of ODE system between bacteria and T cells. For changing initial numbers of bacteria in the range [10^5^, 6×10^7^) at day 0, the system always terminates in stable equilibrium, which biologically corresponds to the chronic infection. Black arrows illustrate the flow of the system. Model parameters are shown in table S1.

We previously demonstrated experimentally that during chronic *S. aureus* infection there is substantial negative correlation between MDSC populations and activated T cells, which are not suppressed (dysfunctional) and hence maintain their ability to proliferate (*6*). To validate the accuracy and consistency of our mathematical model, we reproduced the inverse proportional behaviour between T cells and T-cell suppression by MDSCs (Fig. 1B). Further *in silico* analysis showed how chronic *S. aureus* infection is established naturally, without any treatment intervention (Fig. 1C). Infection induces strong inflammation, which activates T cells. Competition for dominance between bacteria and T cells creates oscillations in the population dynamics (Fig. 1C). To protect from the damaging effect of prolonged inflammation, MDSCs expand gradually to suppress T cells. Increasing accumulation of MDSCs leads to increasing suppression on T cells (Θ). Upon a critical threshold, T-cell suppression by MDSCs is so strong that T cells become dysfunctional and cannot promote sterilizing immunity anymore (Fig. 1D, *pink line*). This *in silico* result agrees with our previous experimental observations, showing that gradual expansion of MDSCs leads to gradual loss of T-cell function, which in turn promotes chronic *S. aureus* infection and failure of sterilizing immunity (*5*, *6*). At this point the immune system finds the balance between maximal bacterial clearance and minimal collateral tissue damage, whereas bacteria persist in the host organism but are simultaneously unable to further grow due to their containment by T cells (*5*) (mathematically known as equilibrium of the system). This equilibrium is found to be stable by our mathematical model (Fig. 1E) and biologically refers to chronic *S. aureus* infection. Once at this stage (Fig. 1D, *pink line;* 1E), sterilizing immunity can be attained only by using treatment-interventions that can destabilize (i.e. perturb) this stable steady state of chronic infection.

### Model-driven therapeutic strategies

To explore perturbation strategies (treatments) that would destabilize the equilibrium between bacteria, T cells and MDSCs, we varied values of *k_b_* and Θ. These parameters, representing T-cell activation and recruitment by bacterial presence, and T-cell suppression by MDSCs, respectively, were specifically chosen because they play a key role in the establishment of chronic infection (*5*, *6*). Different values of *k_b_* and Θ gave different eigenvalues for the ODE system (Eqs. (1)-(2)), which were used to characterize the steady states (equilibria) of the mathematical model as unstable or stable (analytical forms are found in Supplementary Materials). Combining *in silico* all steady states in one separatrix (phase diagram) led to the distinguished areas of cure and chronic infection (Fig. 2A).

**Fig. 2.**
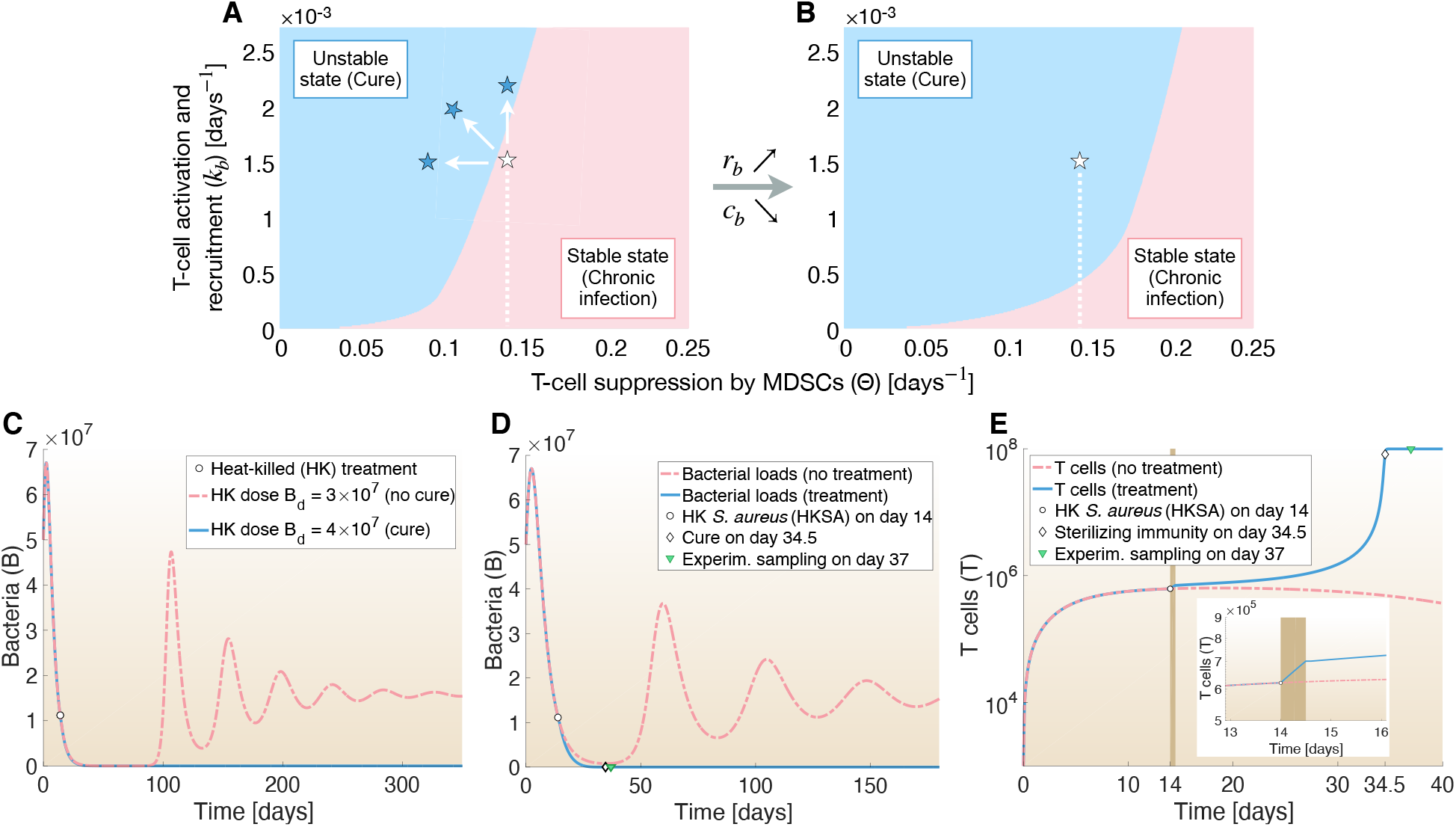
Model-driven insights for cure. (**A-B**) Based on the eigenvalues of the ODE system, the phase diagram was divided into stable states (pink) and unstable states (blue), which represent the physiological chronic infection and cure, respectively. The white star represents the average position of an infected host during a chronic staphylococcal infection and was determined by the fitted values of parameters *k_b_* and Θ (table S1). Given the position of the infected host (white star), cure is achieved (**A**) by increase of T-cell activation and recruitment (*k_b_*) and/or decrease of T-cell suppression by MDSCs (Θ) (shown as blue stars) or (**B**) by increasing the proliferation rate of bacteria (*r_b_*) and/or reducing the bacterial containment by T cells (*c_b_*). (**C-E**) *In silico* treatment with HKSA. Initial infection was induced by setting the bacterial population equal to 5×10^7^ cells at day 0 in Eq. (1). T cells were set to 10^3^ at day 0 in Eq. (2). HKSA-treatment was administered *in silico* at day 14 of infection (shown as white bullet) by adding the term *k_b_B_d_* to the T-cell ODE at time t=perturbation day, for a 12-hour perturbation window as explained in Eq. (3) (*Materials and Methods*). (**C**) A minimum dose of HKSA for curing chronic *S. aureus* infection was estimated to be *B_d_* = 4×10^7^ HKSA. *In silico* administration of 10^8^ HKSA, namely staphylococcal antigens, at day 14 of infection provides (**E**) quick boost of T cells (increase of parameter *k_b_*), as shown in brown, and (**D**) bacterial numbers are diminished to zero. Chronic *S. aureus* infection is cured.

To gain more understanding of the dynamical system and suggest strategies for bacterial clearance, we next estimated the average position of an infected host on the separatrix of cure and chronic infection (Fig. 2A, *white star*), using the values of parameters *k_b_* and Θ as fitted previously (table S1). We found that it lay in the region of chronic infection, yet close to the basin of attraction (region) of cure. According to the separatrix, we concluded that the resolution of chronic infection is achieved via either (a) relocation of the infected host from the chronic infection area towards the area of cure (Fig. 2A) by (i) increasing T-cell activation and recruitment (*k_b_*) and/or by (ii) decreasing T-cell suppression by MDSCs (Θ) or (b) via expansion of the cure zone itself (Fig. 2B) by counter-intuitively (iii) increasing the proliferation rate of bacteria (*r_b_*) and/or (iv) by reducing bacterial containment by T cells (*c_b_*). These *in silico* results accord with previous experimental studies reporting that brief suppression of immunity (namely *c_b_* reduction) with cyclophosphamide (*3*) or targeting MDSCs (namely Θ decrease) (*26*) during *S. aureus* infection reduce bacterial burden. Altogether the model indicates that all four aforementioned perturbation categories (each of which can be implemented *in vivo* in various ways) can destabilize the dynamics of chronic infection so that eradication of *S. aureus* and *sterilizing immunity* is achieved (fig. S1).

### Design of quantitative, model-driven experiments

Our *in silico* analysis suggested treatment strategies that were MDSC-targeting (Θ decrease), pathogen-targeting (*r_b_* increase or *c_b_* decrease), or host-directed (*k_b_* increase). Experimental testing was essential to validate the model predictions. Since validation of bacterial eradication would have immense importance for human therapeutics, we sought to test experimentally a treatment that would be safe and easy to apply.

MDSC-targeting drugs, such as gemcitabine chemotherapeutic agent, usually bear adverse side-effects because of their cytotoxicity. They simultaneously deplete mature immune cells (*26*), which are indispensable components of immunity. Furthermore, available MDSC-targeting drugs fail to eliminate MDSCs in many organs (*27*). Lastly, the heterogeneity of MDSCs makes it particularly difficult to identify a single drug that would deplete all distinct MDSC subsets, each of which is linked to different bacterial infections and even distinct tumors (*8*, *28*).

Contrary to conventional bacteria-targeting drugs, such as antibiotics, that aim at fighting bacteria and whose use should be limited because they promote development of drug-resistance, the bacteria-targeting strategies suggested by our *in silico* analysis counterintuitively promote bacterial growth. However, these strategies would involve treatments, such as immunosuppression (*c_b_* decrease), that would be difficult to implement in reality because, besides cytotoxicity and side-effects that immunosuppressive agents (e.g. cyclophosphamide) cause, patients would be at risk of becoming susceptible to other infections.

Despite ample research focusing on the development of antimicrobial drugs, there is surprisingly a large gap on the development of therapeutics that promote host defense during infection (*29*). Such host-directed therapeutics are safer than pathogen-targeting drugs because they promote the health of the host rather than attack the pathogen (*29*). Therefore, for our experimental testing we singled out the category of *k_b_* increase (namely T-cell activation and recruitment). One of the most conventional and safe ways to boost *k_b_ in vivo*, which is also an established, widely-used method for vaccine development, is via administration of inactivated bacteria, namely antigens. Such treatment would be extremely practical in human therapeutics, since pathogens causing renal abscesses in humans can be isolated from urine samples without involving any invasive processes (*30*) and easily get inactivated. We chose inactivation of bacteria with heat over toxic chemicals, such as formalin. Furthermore, heat-killed cells are more effective, quicker to prepare and provide enhanced survival in treated animals than other inactivation options (*31*). While inactivated bacteria in vaccines serve as prophylaxis from infection, their use as treatment for ongoing chronic infections is limited. Here we explored whether HK treatment during infection leads to cure, as our *in silico* results suggested.

Since our aim was to cure chronic *S. aureus* infection, the experimental perturbation (HK treatment) had to be carried out when infection enters its chronic phase. As we have previously shown, inoculation with 3–7×10^7^ colony-forming units (CFU) of *S. aureus* results in chronic infection and renal abscesses, and by day 14 of infection T cells are already strongly suppressed by MDSCs (*5*, *6*). Therefore, the perturbation with HKSA was scheduled at day 14 after initial infection with 5×10^7^ CFU of *S. aureus*. The physiological *k* increase via HK treatment was incorporated into the model with the addition of term *k_b_B_d_* to the T cells’ ODE on the day of treatment (*Materials and Methods*), where *B_d_* the dose of HKSA and *k_b_* the activation and recruitment of T cells via HKSA assumed the same as for live bacteria during initial inoculation (table S1). This assumption was based on the fact that, although live cells produce virulence factors that stimulate the immune system, implying a greater value of *k_b_*, heat-killed cells cannot hijack or evade immunity as live *S. aureus* does. The only consequence of their existence in the host is the stimulation of immune cells. Additionally, HKSA cells are injected when the host is already chronically infected and had already encountered staphylococcal antigens during initial inoculation. Secondary exposure to pathogens always initiates much stronger (and quicker) immune response than the initial exposure to the same pathogen.

Numerical simulations for initial inoculation with 5×10^7^ *S. aureus* cells and HK treatment at day 14 of infection suggested that the minimum HKSA-dose required for cure would be 4×10^7^ HKSA (Fig. 2C). For our experiments, we opted for the amount of 10^8^ HKSA. To identify, on which day the infected mice would recover from infection to perform the experimental sampling, we followed the *in silico* results, which predicted sterilizing immunity at day 34.5 of infection (Fig. 2D). Because biological systems involve extrinsic and intrinsic stochasticity and therefore not all infected mice are synchronized in the same infection phase, the experimental sampling was set at day 37 of infection (Fig. 2D). As a result of secondary exposure to *S. aureus*, HKSA-treatment initiates a cascade of inflammatory events, providing a concomitant, rapid boost in T-cell population (Fig. 2E, *brown*), which is followed by further, gradual increase of activated T cells as illustrated in our simulations (Fig. 2E). Consequently, HKSA-treatment destabilizes to a sufficient extent the system dynamics of chronic infection and leads to cure (Fig. 2D-2E).

### *In vivo* cure after model-driven perturbation treatment with HKSA

Our next step was to provide proof-of-concept by validating our *in silico* predictions of cure *in vivo*. For this purpose, C57BL/6 mice were infected intravenously with *S. aureus* strain SH1000. At day 14 of infection mice were treated intraperitoneally with HKSA, strain SH1000. Control mice received phosphate-buffered saline (PBS) (Fig. 3A).

**Fig. 3.**
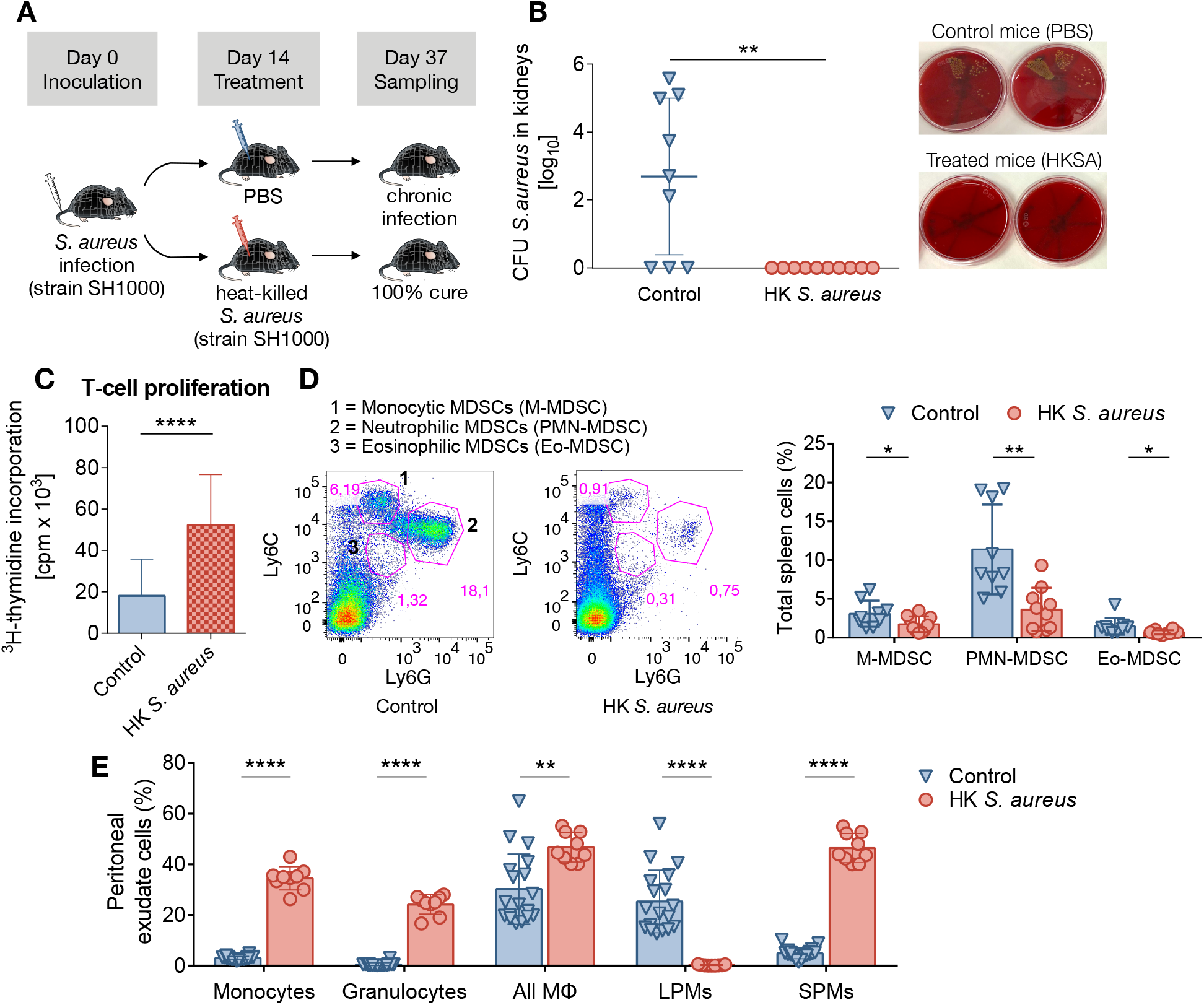
Bacterial burden and immune cells after treatment with heat-killed *S. aureus*. (**A**) Experimental schema. C57BL/6 mice were intravenously inoculated with 5×10^7^ CFU of *S. aureus* and intraperitoneally treated with 10^8^ HKSA at day 14 of infection. Untreated (control) mice received PBS. Sampling was conducted at day 37 of infection. (**B**) Treatment with HKSA at day 14 of infection cures all chronically *S. aureus*-infected mice. Bacterial loads were determined in kidneys of untreated (control) and HKSA-treated mice at day 37 of infection. n=4-5 mice per group, two independent experiments. Data represent mean±SD. **p=0.0018, Student’s t-test. Right: Representative images of ten-fold serial dilutions on blood agar plates for enumeration of viable *S. aureus* (golden-yellow colonies) in kidney homogenates from control (untreated) or HKSA-treated mice, showing high bacterial burden (bacterial persistence) or absence of bacteria (bacterial eradication), respectively. (**C**) Proliferative response of spleen T cells at day 37 of infection from chronically *S. aureus*-infected mice which received PBS or HKSA to *in vitro* stimulation with anti-CD3/anti-CD28. T-cell proliferation was measured by 3H-thymidine incorporation. n=4-5 mice per group, two independent experiments. Data represent mean counts per minute (cpm) ± SD. ****p<0.0001, Student’s t-test. (**D**) HKSA-treatment reduces MDSCs. (left) Representative fluorescence-activated cell sorting (FACS) illustrating monocytic CD11b^+^Ly6C^+^Ly6G^low^, neutrophilic CD11b^+^Ly6C^low^Ly6G^+^ and eosinophilic CD11b^+^Ly6C^low^Ly6G^low^ MDSCs, and (right) percentage of each MDSC subset within the total spleen cell population of untreated (control) or HKSA-treated mice at day 37 of infection. n=4-5 mice per group, two independent experiments. Data represent mean±SD. *p<0.05 and **p<0.01, Student’s t-test. (**E**) Intraperitoneal HKSA-treatment at day 14 of infection induces acute inflammation. Peritoneal exudate cells were collected from chronically *S. aureus*-infected mice 12 hours after administration of PBS (control) or HKSA. Percentage of CD11b^+^Ly6C^+^ monocytes, CD11b^+^Ly6G+ granulocytes, CD11b^hi^F4/80hi LPMs, CD11b^+^F4/80^low^ SPMs along with total MФ population (LPMs plus SPMs) within the total peritoneal cell population is shown. n=5-10 mice per group, two independent experiments. Data represent mean±SD. **p<0.01, ****p<0.0001, Student’s t-test.

Our previous study in chronically *S. aureus* SH1000-infected C57BL/6 mice has shown that *S. aureus* is progressively depleted from multiple organs and persists only in the kidneys (*5*). Therefore, at day 37 of infection, bacterial load quantification was performed in mice’s kidneys. The mathematical model’s predictions *in silico* (Fig. 2D) were validated by our experiments *in vivo*: mice treated with HKSA were cured with success percentage of 100% and no *S. aureus* at all was found in their kidneys (Fig. 3B). In contrast, the majority of untreated animals remained infected with high bacterial burden (Fig. 3B).

Progression of *S. aureus* infection from acute to chronic drives spleen T cells into T-cell dysfunction, which we previously found irreversible (*5*). T-cell dysfunction during chronic *S. aureus* infection is caused by the expansion and suppressive effect of all three MDSC subsets in spleen (*6*, *13*). Therefore, at day 37 of infection, we next investigated the effect of the HKSA-treatment on the proliferative response of T cells and on the MDSC populations. We found that HKSA-treatment restored the function of spleen T cells. Stimulation with anti-CD3/anti-CD28 antibodies showed that T cells were hyperresponsive and actively proliferated (Fig. 3C). In contrast, spleen T cells of infected untreated animals remained hyporesponsive to TCR re-stimulation (Fig. 3C). We further observed significant reduction of all MDSC subsets in spleens of HKSA-treated mice, whereas MDSC populations remained high in spleens of infected, untreated (control) animals, which had received PBS (Fig. 3D). Together, these findings reveal that HKSA-treatment during chronic *S. aureus* infection not only boosts T-cell function (namely *in vivo k_b_* increase) but also has a previously unidentified potential to indirectly target the MDSCs (namely *in vivo* Θ decrease). Because MDSC-expansion and MDSC-mediated suppression on T cells are associated with the chronic but not acute phase of the infection (*5*, *6*, *8*), these results suggest that acute inflammation caused by administration of HKSA (see below) disrupts the balance of chronic infection, which sustains MDSCs, leading to natural MDSC-depletion.

### HKSA-treatment during chronic *S. aureus* infection induces strong acute inflammation

Our *in silico* analysis suggested that destabilizing the system of chronic *S. aureus* infection by administering a sufficient amount of HKSA would lead to cure. It was natural to expect that the insertion of staphylococcal antigens into the hosts via HKSA-injection would trigger acute inflammation. To verify *in vivo* that the HKSA-injection initiated acute inflammation during chronic *S. aureus* infection, we sampled peritoneal exudates 12 hours after intraperitoneal HKSA-treatment and assessed differences in populations of macrophages, as well as infiltrating monocytes and granulocytes compared to untreated mice. It has been previously shown that two distinct subsets of macrophages exist in mouse peritoneal cavity (PerC), the CD11b^hi^F4/80^hi^ large peritoneal macrophages (LPMs) and the CD11b^+^F4/80^low^ small peritoneal macrophages (SPMs), and together are responsible for most of phagocytosis happening in PerC (*32*, *33*). Under normal physiological conditions, LPMs are the predominant macrophage subset in PerC (*32*). However, under inflammatory conditions, the PerC environment changes drastically: LPMs disappear and SPMs become the major subset along with substantial recruitment of SPM-precursors, the CD11b^+^Ly6C^+^ monocytes (*32*) and of CD11b^+^Ly6G+ granulocytes (neutrophils). In accordance, we observed massive increase in amounts of CD11b^+^Ly6C^+^ monocytes and CD11b^+^Ly6G+ granulocytes (neutrophils) in PerC of HKSA-treated mice compared to untreated (control) mice (Fig. 3E). LPMs were the predominant macrophage subset in chronically *S. aureus*-infected, untreated mice, indicating homeostatic conditions in PerC, whereas treatment of chronically *S. aureus*-infected mice with HKSA resulted in disappearance of LPMs and predominance of SPMs, indicating acute inflammation (Fig. 3E). Total number of MФ (LPMs plus SPMs) was increased in PerC of HKSA-treated mice (Fig. 3E). These results suggest that treatment with HKSA during chronic *S. aureus* infection induced strong acute inflammatory responses.

### Non-antigen-specific HK treatment

The mathematical model suggested that treatment with HK *S. aureus* would cure chronic *S. aureus* infection. Because MDSCs during chronic *S. aureus* suppress T cells, including antigen-specific T cells, we proceeded to assess experimentally whether the HK treatment (perturbation strategy) works in an antigen-specific manner. For reliable comparison with the experiments using HKSA, we maintained the experimental design of the aforementioned quantified HKSA-protocols (namely same CFU of *S. aureus* for inoculation, HK-dose, routes of administration, times of treatment and sampling) and only replaced HKSA with HK cells of a different bacterium. Therefore mice were inoculated with 5×10^7^ *S. aureus*, however, they were treated intraperitoneally with 10^8^ HK *Streptococcus pyogenes* at day 14 of infection. Sampling was performed at day 37 of infection (Fig. 4A). HKSP-treatment successfully cured 50% of chronically *S. aureus*-infected mice (Fig. 4B), whereas the majority of control mice remained infected. Similarly to experiments with HKSA-treatment, we explored how HKSP-treatment affected T-cell proliferation and MDSC populations. Spleen T cells of HKSP-treated mice responded to stimulation with anti-CD3 plus anti-CD28 antibodies and proliferated significantly more than spleen T cells of untreated mice, which remained hyporesponsive (Fig. 4C). In contrast to HKSA-treatment that led to considerable MDSC-depletion, the reduction of MDSCs in HKSP-treated mice was insignificant compared to MDSCs in untreated mice (Fig. 4D).

**Fig.4.**
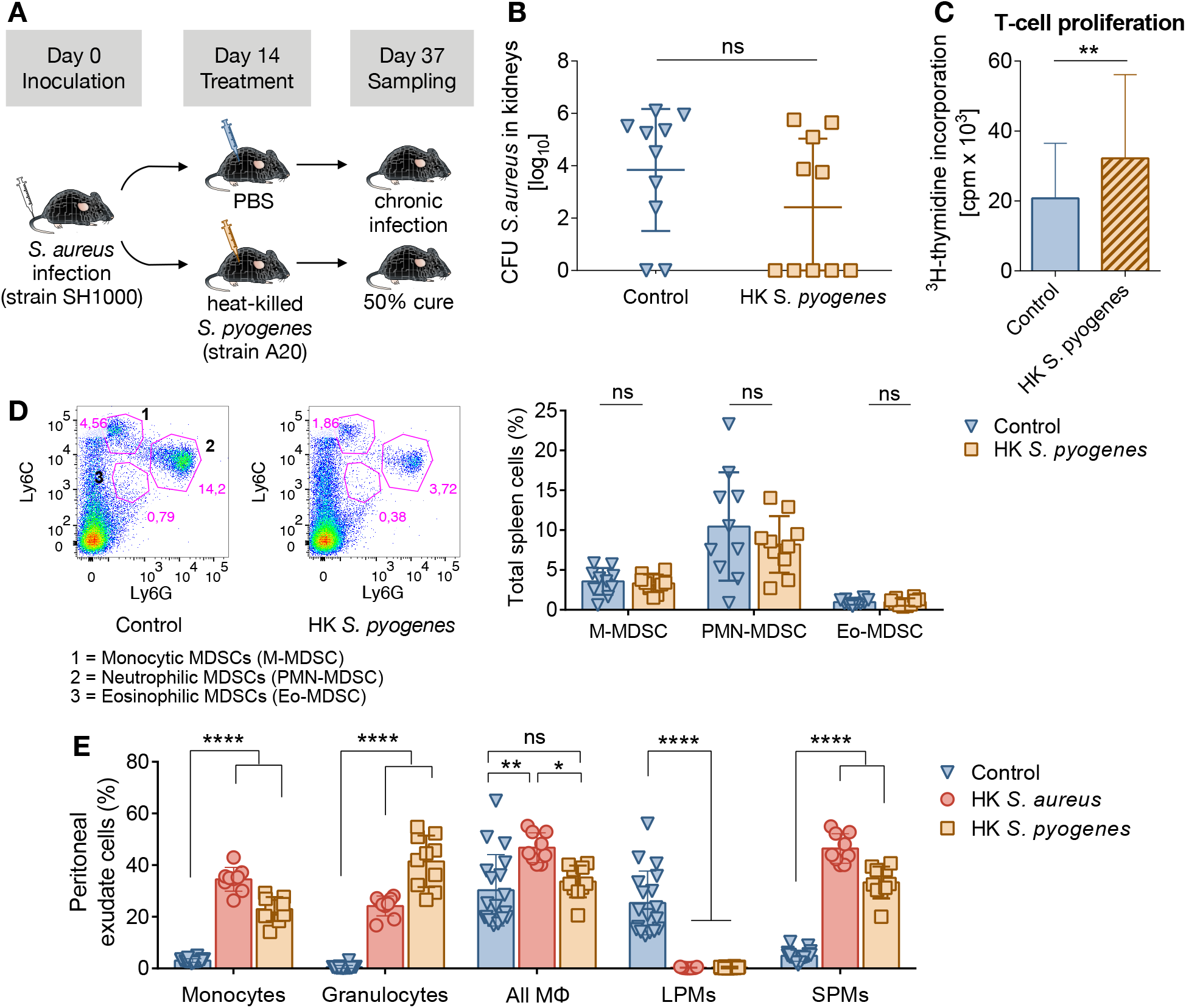
Bacterial burden and immune cells after treatment with heat-killed *S. pyogenes*. (**A**) Experimental schema. C57BL/6 mice were intravenously inoculated with 5×10^7^ CFU of *S. aureus* and intraperitoneally treated with 10^8^ HKSP at day 14 of infection. Untreated (control) mice received PBS. Sampling was conducted at day 37 of infection. (**B**) Treatment with HKSP at day 14 of infection cures 50% of chronically S. aureus-infected mice. Bacterial loads were determined in kidneys of untreated (control) and HKSP-treated mice at day 37 of infection. n=5 mice per group, two independent experiments. Data represent mean±SD, Student’s t-test. (**C**) Proliferative response of spleen T cells at day 37 of infection from chronically *S. aureus*-infected mice which received PBS or HKSP to *in vitro* stimulation with anti-CD3/anti-CD28. T-cell proliferation was measured by ^3^H-thymidine incorporation. n=5 mice per group, two independent experiments. Data represent mean counts per minute (cpm) ± SD. **p=0.0024, Student’s t-test. (**D**) HKSP-treatment slightly reduces MDSCs in spleen. Representative fluorescence-activated cell sorting (FACS) illustrating monocytic CD11b^+^Ly6C^+^Ly6G^low^, neutrophilic CD11b^+^Ly6C^low^Ly6G^+^ and eosinophilic CD11b^+^Ly6C^low^Ly6G^low^ MDSCs (left), and percentage of each MDSC subset within the total spleen cell population of untreated (control) or HKSP-treated mice (right) at day 37 of infection. n=5 mice per group, two independent experiments. Data represent mean±SD, Student’s t-test. (**E**) Intraperitoneal injection with HKSP at day 14 of infection induces acute inflammation. Peritoneal exudate cells were collected from chronically *S. aureus*-infected mice 12 hours after administration of PBS (control) or HKSP. Percentage of CD11b^+^Ly6C^+^ monocytes, CD11b^+^Ly6G^+^ granulocytes, CD11b^hi^F4/80^hi^ LPMs, CD11b^+^F4/80^low^ SPMs along with total MΦ population (LPMs plus SPMs) within the total peritoneal cell population is shown. n=5-10 mice per group, two independent experiments. Data represent mean±SD. *p<0.05, **p<0.01, ****p<0.0001, one-way ANOVA with Tukey’s multiple comparisons test. Peritoneal exudate cells collected 12 hours after HKSA-treatment at day 14 of infection are included for comparison.

As in the case of HKSA-treatment, peritoneal exudates were sampled 12 hours after HKSP-treatment and confirmed acute inflammation. CD11b^+^Ly6C^+^ monocytes and CD11b^+^Ly6G^+^ granulocytes (neutrophils) increased massively in PerC of HKSP-treated mice compared to untreated mice (Fig. 4E). LPM-disappearance further confirmed acute inflammation induced by HKSP-treatment along with significant increase of SPMs (Fig. 4E). It is important to note that after HKSA-treatment, which cured all infected hosts, CD11b^+^Ly6C^+^ monocytes increased massively, whereas after HKSP-treatment, which cured half of the infected hosts, CD11b^+^ Ly6G^+^ granulocytes (neutrophils) increased massively (Fig. 4E).

These results confirmed that HK treatment with either *S. aureus* or *S. pyogenes* induces acute inflammation during chronic *S. aureus* infection. However, it was expected that HKSA-treatment would initiate stronger acute inflammation than HKSP-treatment, because HKSA was a re-exposure to *S. aureus*, while HKSP-treatment was a first-time exposure to *S. pyogenes* antigens. This is reflected by a significant increase of total MΦ (LPMs plus SPMs) in PerC after HKSA but not after HKSP-treatment compared to control, untreated mice (Fig. 4E). Weaker acute inflammation after HKSP-than HKSA-treatment was also indicated by less reduction of MDSCs in spleens (Fig. 3D, 4D), which are sustained in conditions of chronic (but not acute) inflammation (*8*) in addition to almost twofold lower proliferative response of spleen T cells in HKSP-treated compared to HKSA-treated mice (Fig. 3C, 4C). These results combined can explain cure in 50% of HKSP-treated animals in comparison to cure in 100% of HKSA-treated animals and accord with previous studies, suggesting that monocytes and macrophages promote *S. aureus* clearance (*26*) and that after antigen immunization in PerC, SPMs migrate to lymph nodes where they activate T cells (*33*, *34*).

Because treatment with 10^8^ HKSP induced weaker acute inflammation and T-cell stimulation than treatment with 10^8^ HKSA, we hypothesized that treatment with higher HKSP-dose (more antigens) would naturally induce stronger acute inflammation (including more CD11b^+^Ly6C^+^ monocytes and SPMs), hence more indirect depletion of MDSCs and more activated T cells, leading to cure of all HKSP-treated mice. For this purpose, we reduced the value of parameter *k_b_* in the treatment term *k_b_B_d_* by two or three times to reflect the weaker T-cell stimulation by HKSP compared to HKSA and estimated day of clearance for varying HKSP-doses *B_d_* administered at day 14 of infection. Our simulations suggested that if higher HKSP-dose (>10^8^) is administered, all HKSP-treated mice could be cured (namely 100% success) by sampling day, day 37 of infection (fig. S2).

As commonly known, secondary exposure to antigens elicits a faster immune response, meaning that HKSA-treatment, which was a re-exposure to staphylococcal antigens, triggered a much faster immune response compared to HKSP-treatment, which was a first-time exposure to streptococcal antigens. Therefore, for HKSP-dose the same as HKSA-dose, HKSP-treatment would require longer time to induce immune response relatively as strong as that induced by HKSA. Consequently, although the cure of 50% of HKSP-treated mice at day 37 of infection, when experimental sampling was conducted, was not statistically significant (Fig. 4B), we hypothesized that at a later time-point HKSP-treatment could eventually cure all hosts. For this purpose, we reduced the value of parameter *k_b_* in the treatment term *k_b_B_d_* by two or three times to reflect the weaker T-cell stimulation by HKSP compared to HKSA and we estimated the day of clearance for 10^8^ HKSP (*B*) administered at day 14 of infection. Indeed, our simulations suggested that if sampling is conducted approximately 10 days or later than day 37 of infection, all HKSP-treated mice may be cured, namely 100% success (fig. S2).

### Reasoning for past unsuccessful applications of the treatment

Our model-driven protocol suggesting sufficient HKSA-dose, its administration day, and day of complete bacterial clearance has proved reliable and effective for all HKSA-treated animals. Although administration of inactivated cells as treatment for infections has been used in the past, this kind of therapy is narrowly established. This is due to lacking information regarding the sufficient dose(s) required and day(s) of administration that could resolve *S. aureus* infections successfully. Until now, treatments with inactivated bacteria against *S. aureus* infections have been based on vague experimental experience.

Here, we explain *in silico* why heat- or formalin-killed bacteria treatments used so far have not been successful in yielding clearance. We base our arguments on a previous study (*35*), in which scientists administered at least 19 injections with increased dose of formalin-killed *S. aureus* over the period of 3 months in human patients with *S. aureus* infection (furunculosis).

In the study none of the chronically infected patients was reported to have attained sterilizing immunity, even though they experienced moderate to strong clinical improvement. The injecting scheme in the study consisted of increasing killed *S. aureus* doses (*B_d_*) given in intervals of 3-5 days as follows:

– Suspension I:(0.1,0.2,0.3,0.4,0.5)×5×10^8^
– Suspension II: (0.3, 0.4, 0.5, 0.6, 0.7, 0.8, 0.9, 1)×10^9^
– Suspension III:(0.5,0.6,0.7,0.8,0.9,1)×2.5×10^9^.

We assumed that the bacterial capacity in humans is 1000 times greater than the bacterial capacity in mice (based on the kidney weight ratio between mice and humans), created the corresponding murine suspensions:

– Suspension IV:(0.1,0.2,0.3,0.4,0.5)×5×10^5^
– Suspension V: (0.3, 0.4, 0.5, 0.6, 0.7, 0.8, 0.9, 1)×10^6^
– Suspension VI:(0.5,0.6,0.7,0.8,0.9,1)×2.5×10^6^,

and applied them in our mouse model *in silico* every 4 days, starting from day 14, when chronic infection is established in mice.

The conventional administration of injections with inactivated bacteria is based on the belief that repeated vaccination could work more efficiently. Here, we employed *in silico* the analogous protocol that has been administered in humans (suspensions IV-VI, 19 injections) and showed that repeated injections with increasing doses of killed *S. aureus* cannot render eradication of *S. aureus* (Fig. 5A). We also explored the case when not only more injections are given (25 in total) but also with high fixed dose (almost fivefold higher than the highest dose of suspensions IV-VI) and found that such treatments still fail to eliminate the infection (Fig. 5A). Our results explicitly negate the current belief by showing that challenging the system of chronic infection repeatedly with insufficient killed *S. aureus* dose and in high-frequency intervals does not result in cure (Fig. 5A).

**Fig. 5.**
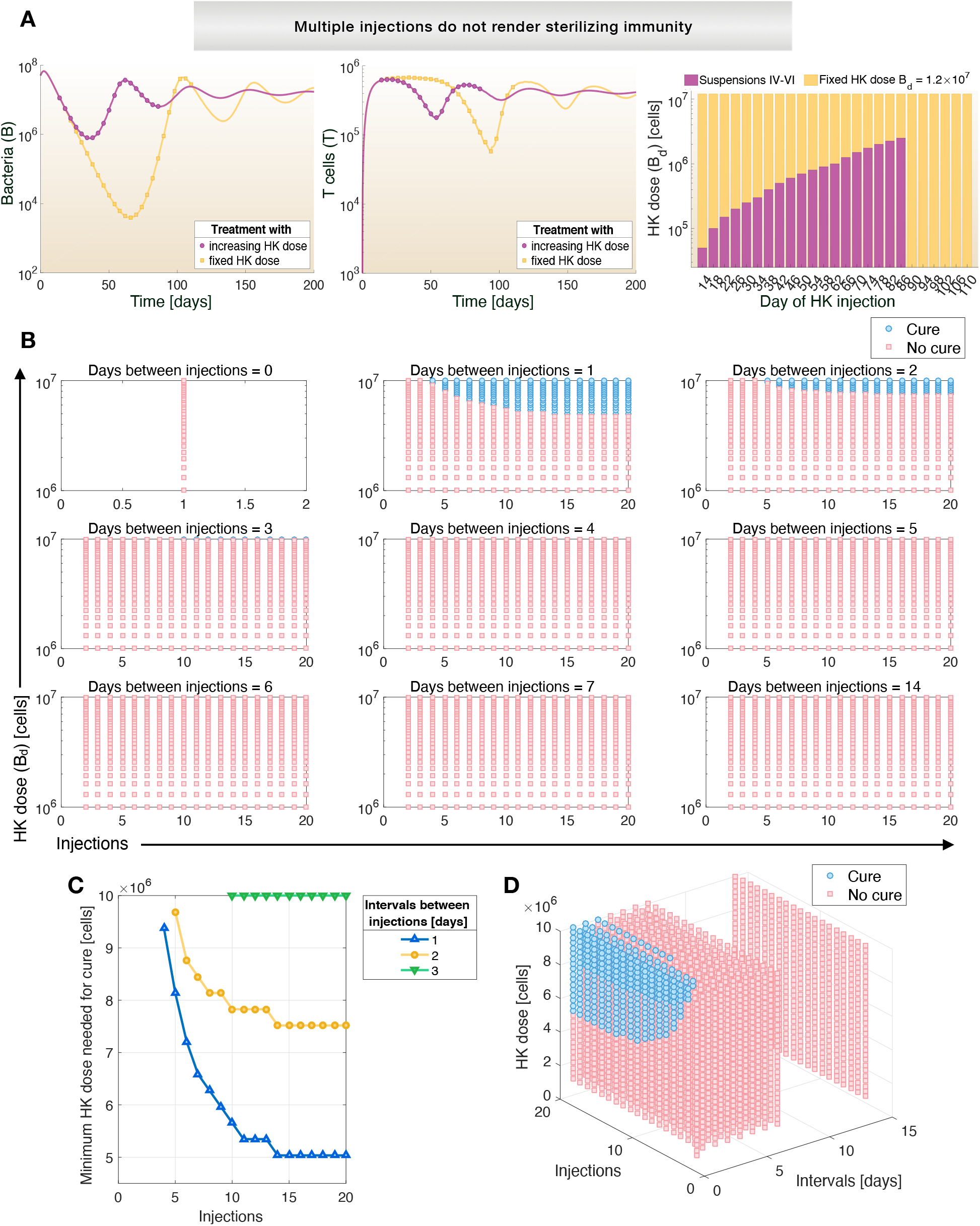
Simulations explain why previous treatments with inactivated *S. aureus* in humans were unsuccessful and suggest curative treatment protocols. At each injection day starting from day 14 of infection (when infection is chronic), treatment was incorporated into the T-cell ODE (Eq. (2)) as *k_b_B_d_*, where *k_b_* as defined in table S1 and *B_d_* the varying, administered HKSA-dose. (**A**) Repeated administration of inactivated *S. aureus* cannot render clearance. 19 HKSA-injections with increasing dose in an analogous way to as administered in humans (suspensions IV-VI, purple) or 25 injections with high fixed dose (yellow) were administered every four days. Chronic *S. aureus* infection persists. (**B-D**) *In silico* predictions for cure plotted for 1 up to 20 HKSA-injections, HKSA-doses ranging from 10^6^ to 10^7^, and time intervals between HKSA-injections from 0-7 days, or 2 weeks. Time interval of 0 days means that only one HKSA-injection is administered. (**B**) Possibility of cure declines when time intervals between HK-injections increase, even if many injections with high HKSA-dose are administered. Each blue circle represents a unique, curative treatment protocol and shows that cure is achieved only when time intervals between injections are 1-3 days with minimum HKSA-dose needed for cure as in (**C**). (**D**) Interconnection between HKSA-doses 10^6^ to 10^7^, HKSA-injections and time intervals between injections in three-dimensional plot.

To gain a broader perspective on curative protocols, we simulated varying doses from 10^6^ to 10^7^ inactivated *S. aureus*, number of injections from 1-20 and time intervals between injections from 0 up to 14 days (Fig. 5B-5D). Our *in silico* results suggested that for doses of 10^6^ to 10^7^ inactivated *S. aureus,* only treatment protocols involving HK injections with time intervals from 1-3 days could cure (Fig. 5B, *blue*), with the maximum time interval of 3 days requiring a minimum dose of 10^7^ inactivated *S. aureus* to be effective (Fig. 5C). Therefore, both the time intervals of 3-5 days and doses (suspensions IV-VI) analogous to as administered to furunculosis patients (*35*) were fruitless attempts towards bacterial clearance, since the administered doses in those time intervals were low to sufficiently stimulate the immune system and lead to cure (Fig. 5B-5D). We next postulated that higher doses could have cured chronically *S. aureus*-infected patients. For this purpose, we simulated curative and non-curative protocols for varying number of injections, time intervals between injections and doses from 8 × 10^6^ up to 10^8^ inactivated *S. aureus*. Our results clearly indicated that higher doses could lead to cure (Fig. 6A-6C). Furthermore, our results showed that in case time intervals between injections need to increase, (e.g. schedules of doctors), then higher HK-doses are required for cure (Fig. 6B).

**Fig. 6.**
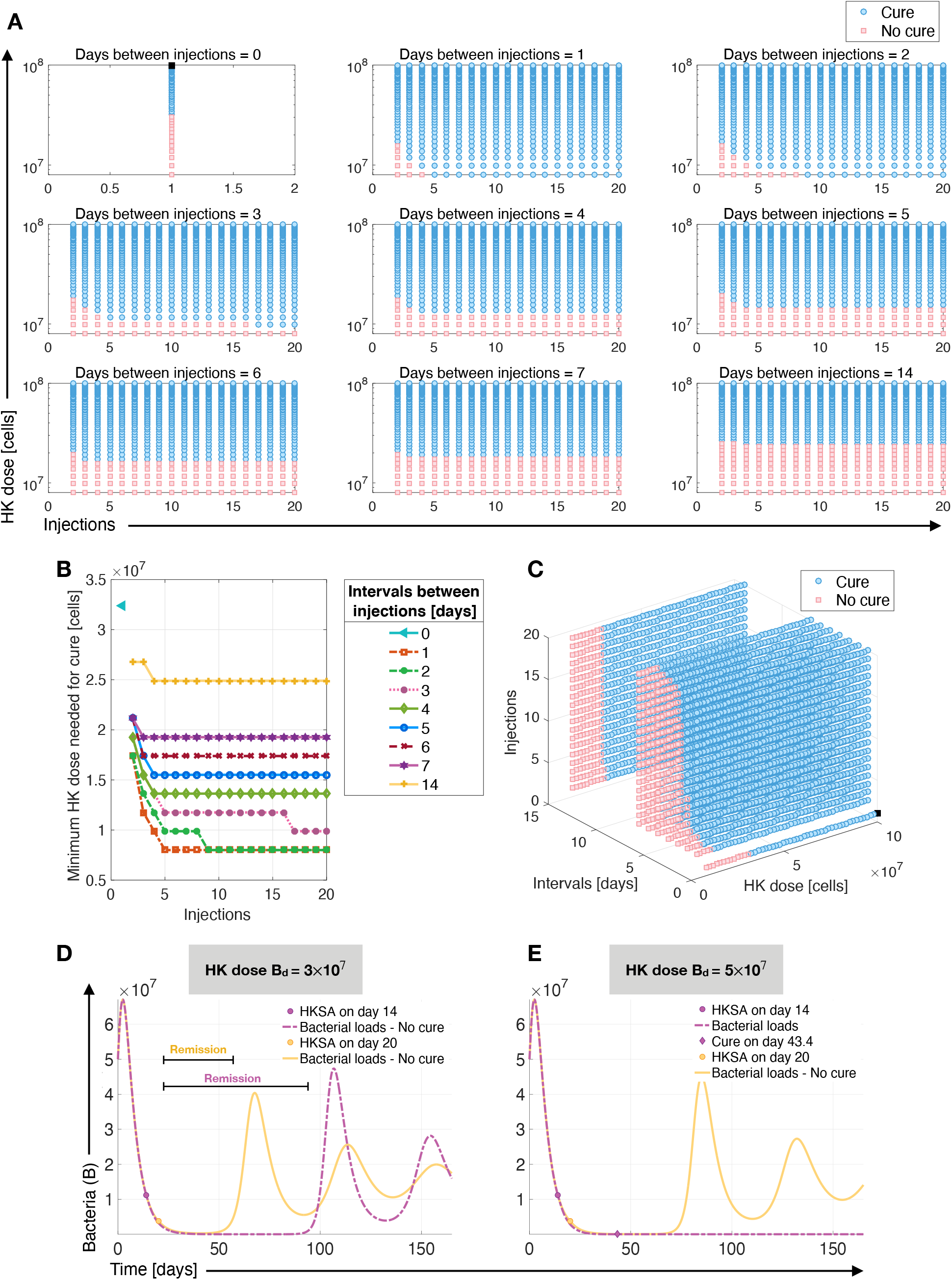
*In silico* correlation between HKSA-doses from 8×10^6^ to 10^8^, number of HKSA-injections and time intervals between injections for curative treatment protocols or remission. (**A-C**) Same as in Fig. 5B-5D for HKSA-doses ranging from 8×10^6^ to 10^8^. Each blue circle represents a unique, curative treatment protocol, for which the minimum dose required for successful treatment is indicated in (**B**). Black square in (**A**) and (**C**) represents the treatment protocol of 10^8^ HKSA in a single injection that was tested experimentally in this study and cured all hosts *in vivo*. (**D-E**) Influence of administration time of HKSA-injection on the chronic infection. The HKSA-treatment *in silico* was added as term *k_b_B_d_* to the T-cell ODE (Eq. (2)) at day 14 or 20 for a 12-hour perturbation window, where *k_b_* as in table S1 and *B_d_* the HKSA-dose. Critical HKSA-dose required for cure when administered at day 14 (early chronic infection) was estimated to be 4×10^7^ (Fig. 2C). Although lower HKSA-dose (**D**) for example *B*_d_ =3×10^7^ HKSA cannot cure, it provides remission. Higher HKSA-dose (**E**), such as *B*_d_ =5×10^7^, confers sterilizing immunity when administered at early timepoints of chronic infection but fails to cure when administered later, although it provides remission temporarily.

Even though treatments with insufficient-for-cure doses of inactivated *S. aureus*, such as the doses given to furunculosis patients, cannot confer complete clearance of bacteria, furunculosis patients still reported improvement. In accordance, *in silico* administration of low HK-doses showed that although they do not cure the hosts, they alleviate chronic infection by providing remission (Fig. 6D). In fact, administration of HK treatment as early as possible leads to longer remission of the infection (Fig. 6D). Lastly, we investigated how the administration time of a HK-injection affects the clinical outcome. For middle-sized doses the day of administration is crucial for the outcome of chronic infection, since it can provide cure if given as early as possible or only provide temporary remission without clearing the infection (Fig. 6E). These results highlight the urgency of treating chronic *S. aureus* infections as early as possible, which is often compromised in cases of *S. aureus*-caused renal abscesses by improper and/or time-consuming treatments (*16*, *30*).

## DISCUSSION

At present, no treatment has proved to be completely effective in resolving chronic *S. aureus* infections. There is no vaccine against *S. aureus* infections because all clinical trials have failed (*36*). The development of new antibiotics would only be a temporary solution, until the bacterium develops antibiotic resistance. Recent reports warn on the alarming increase of antibiotic-resistant *S. aureus* infections in organs that have not been conventionally infected before, including kidneys (*16*, *37*). Such reports highlight that treating infections in organs, where the medical community is not traditionally used to encountering, is very challenging and often misdiagnosed, therefore requiring prompt intervention once identified (*30*). Furthermore, the current medical arsenal of antibiotics, MDSC-targeting drugs or surgery, is insufficient, ineffective or inadvisable due to bacteria-resistance mechanisms, cytotoxicity of drugs, inability to drain abscesses because of their location in internal organs or the high-risk medical status of patients. Consequently, new, less complicated ways of treatment against *S. aureus* infections are absolutely essential to find. We here reported cure of all chronically *S. aureus*-infected mice with renal abscesses, using heat-killed *S. aureus* treatment, as quantified by our mathematical model. Although inactivated *S. aureus* has been used to treat soft skin infections *(35)* this is, to our knowledge, the first time heat-killed *S. aureus* is used to treat renal abscesses.

Conventional vaccination with inactivated microorganisms always requires multiple doses to ensure prophylaxis. Hence, it is believed that inactivated microorganisms, when used as treatment, should also be administered in multiple doses to be efficient (*35*). Here we provided evidence that treatment with one HK-injection alone can cure chronic *S. aureus* infection (Fig. 2D, 3B). This is an important finding, because it means that a single injection with the right HK-dose, besides being extremely practical to implement in human therapeutics, can suffice to break the chronic phase of infection and lead to bacterial clearance, without the need for antibiotics, that multidrug-resistant *S. aureus* can evade, or surgery for abscess drainage, or cytotoxic MDSC-targeting drugs. Most importantly, pathogens causing renal abscesses in humans are easily obtained without any invasive methods from urine samples from patients (*30*), which can then be easily cultured and heat-killed. Regarding safety, the administration of inactivated *S. aureus* as treatment of staphylococcal skin infection in human patients had only minor side-effects, such as temporary pain at the site of injection (*35*). In accordance, our treatments with HKSA or HKSP were well-tolerated. Treated animals did not exhibit any signs of side-effects or discomfort. They were active, alert, observant of their environment, with eyes wide open, fur laying flat and maintained normal body weight and appetite. Consequently, HK treatments bear the potential to become easily produced, easily stored, cost-effective, and safe treatments in medicine.

The recently identified Eo-MDSC inhibit, as other MDSCs, T-cell proliferation during chronic *S. aureus* infection (*13*). Our results agree with previous results on the suppressive effect of Eo-MDSC on T cells, since high Eo-MDSC numbers in control (untreated) mice contributed to high T-cell suppression that was reduced as a result of reduced Eo-MDSC after HKSA-treatment (Fig. 3C-3D). More Eo-MDSC have been directly linked with more CFU of *S. aureus* (*13*). In accordance, our results showed that significant decrease of Eo-MDSC contributed to bacterial clearance (Fig. 3B, 3D). Additionally, we here revealed a previously unidentified approach for depletion of Eo-MDSC that was achieved in an indirect way, in contrast to cytotoxic agents, which directly target MDSC-populations and bear side-effects.

Our mathematical model aimed to predict variables that could transition a chronic *S. aureus* infection to an active inflammatory state to promote bacterial clearance. Our experiments *in vivo* verified that perturbation with HKSA initiates acute inflammation during the chronic establishment of *S. aureus* infection, showing massive increase in amounts of CD11b^+^Ly6C^+^ monocytes, CD11b^+^Ly6G^+^ granulocytes (neutrophils) and CD11b^+^F4/80^low^ SPMs in HKSA-treated mice (Fig. 3E). Our previous study has shown that T cells from chronically *S. aureus*-infected mice remained hyporesponsive after *in vitro* stimulation with either heat-killed *S. aureus* or anti-CD3 plus anti-CD28 (*5*). This indicates that it is not the heat-killed treatment per se that cured the infection, but rather the mathematically-driven concept of sufficiently destabilizing the system dynamics *in vivo*. One of the strategies for destabilization of the chronic infection system, as shown in this study, was achieved via administration of HKSA that caused acute inflammation and subsequently created a new cascade of inflammatory events (including T-cell boost), resulting in sterilizing immunity.

For the experiments regarding antigen-specificity of the treatment, *S. pyogenes* was chosen because together with *S. aureus* they are the two most common gram-positive cocci of medical significance (*38*). Delivery of 10^8^ HKSP cleared *S. aureus* in 50% of animals, suggesting that a portion of the effects of the HK treatment are antigen-independent and that a non-antigen specific HK treatment may be completely effective if combined with other perturbation strategies (antigen-specific or not). Even a higher HKSP-dose alone could have potentially resolved the chronic infection in all HKSP-treated animals (fig. S2). Interestingly, we found that other studies have used a similar approach to treat chronic *Mycobacterium tuberculosis* infections with heat-killed *Mycobacterium vaccae,* especially in patients, in whom previous treatment with antibiotics and/or chemotherapy had failed. Consistent with our results, they also reported a therapeutic effect that cures patients (success percentages varying), along with increase of T cells and an exemplary safety record even in patients co-infected with human immunodeficiency virus (HIV). The copious advantages of heat-killed *M. vaccae* treatments led to extensive Phase III clinical trials in patients in a plethora of countries around the globe (*39, 40*). Taking all the above into account, our results with *S. pyogenes* could be pioneer for chronic *S. aureus* infections because they suggest that chronic *S. aureus* infections may be successfully treated with other *S. aureus* strains or even other bacteria, for example *S. pyogenes*. If the presented results are corroborated by further evidence and hold in human, a standardized cocktail of the most frequent organisms causing renal abscesses can be added to hospital stocks for immediate treatment of patients afflicted with renal abscesses. In addition, these results could also help current efforts towards vaccine development for protection against *S. aureus*, since antigens from other cocci/bacteria could be incorporated for vaccine testing.

Some diversity in the bacterial loads of control mice in all experiments *in vivo* was observed (Fig. 3B, 4B). All control mice were infected with *S. aureus* but did not receive heat-killed treatment. It was observed that in some of them *S. aureus* was cleared. Such variability can be occasionally observed since (i) individual immune responses of mice can vary greatly and (ii) biological systems are inherently stochastic. However, chronic infection and associated renal abscesses caused by the same *S. aureus* strain as used here have been amply studied and the dynamics of chronicity *in vivo* were published already years ago (*5*, *6*). Even though these controls would be typically discarded, we included them for reasons of scientific transparency and because this observed behaviour is explained mathematically in Fig. 2A: The average position of an infected animal in the separatrix (*white star*) lies in the basin of attraction of chronic infection (Fig. 2A, *pink area*) but is very close to the basin of attraction of cure (Fig. 2A, *blue area*), indicating that a small portion of infected mice could spontaneously get cured. Mathematically, points that lie close to the boundary separating two basins of attraction could end up in the other attractor domain, namely the cure region, after stochastic perturbations (possibly happening during the acute phase of the infection). Most essential is, however, the effectiveness of the treatment because renal abscesses are often misdiagnosed, hence associated with high risk of mortality (*30*), and once correct diagnosis takes place, the patients might already experience organ damage and be in life-threatening conditions, whose lives could be saved because of this treatment.

Our mathematical model is not limited to our proof-of-concept presented here. The potential, novelty and usefulness of our mathematical model are further depicted in Fig. 6, showing a plethora of unique, curative HKSA-treatment protocols with varying HK-doses, number of HK-injections and time intervals between HK-injections (each blue bullet point). This variety of quantified protocols can offer important flexibility for future treatments or clinical trials because they can comply with the scientific and social circumstances, for instance working hours of doctors, sensitivity of individuals to high HK-doses, or restrictions in animal permits. We have started designing a graphical user interface for all these protocols, so that scientists can choose HK-dose(s) or time intervals between treatment(s) and easily design their research and treatments.

The results of this study are specified to C57BL/6 female mice; however, new *in silico* curative protocols can be obtained when fitting the parameters of the mouse line or human of interest. Most importantly, the simple structure of our model allows its easy adjustment for chronic infections caused by other bacteria or staphylococcal strains, which could suggest curative protocols against other strain- or bacteria-specific infections, including *S. aureus* biofilms on medical implants that are devastating the clinics, renal abscesses caused by other bacteria such as *Escherichia coli*, or infections in other internal organs. Future studies could be such adjustment of model parameters for other bacteria or staphylococcal strains and subsequent experiments based on model-driven treatment protocols for verification.

We here report cure of chronic *S. aureus* infection and renal abscesses without any use of antibiotics, cytotoxic MDSC-targeting drugs, or surgical procedures for abscess drainage. Our study provides proof-of-principle for a novel treatment protocol of chronic *S. aureus* infection by repurposing heat-killed treatments, guided and quantified by our mathematical model. Unlike conventional heat-killed administration, which is used as prophylaxis against infections in humans and sometimes animals, we used this method as a treatment during ongoing chronic infection. Treatments that are directed to promoting the immune response during an ongoing infection are lacking. Our work could pave the way and provide a new scientific perspective towards treatments that do not aim at targeting the infective agent (such as antibiotics), but rather boost the host’s own immune defense to resolve the infection.

## METHODS

### Mathematical model

The mathematical model was implemented and simulated in MATLAB, see www.mathworks.com.

#### Simulating the perturbation treatment

To simulate the perturbation strategy, we incorporated, for a perturbation window of 12 hours, the extra term *k B* into the ODE describing T cells (Eq. (2)), where *B* = 10^8^ cells is the dose of heat-killed *S. aureus* and *k*_b_ = 0.001509 [days^−1^] the parameter of T-cell activation and recruitment as estimated during the fitting process (full description in Supplementary Materials; table S1). The term of heat-killed treatment was incorporated into the ODE of T cells, because heat-killed cells, as it happens when administered via conventional vaccines, deliver antigens into the body that enhance the adaptive immunity. On each day (timepoint) of administration of heat-killed treatment, such as day 14 of infection, the treatment term *k_b_B_d_* was added, and the equations became:

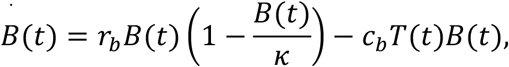

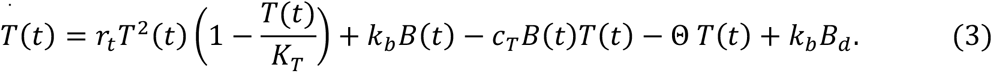

Since heat-killed bacteria are not present continuously *in vivo*, the treatment term (that simulates the *in vivo* conditions) was removed after the perturbation window and the equations returned to their initial form as in equations (1)-(2). For treatment with heat-killed *Streptococcus pyogenes* or varying HK-dose, the HK-dose *B_d_* and value of *k_b_* in the treatment term *k_b_B_d_* were as stated in the main text.

### Experimental protocols

#### Bacteria

*S. aureus* strain SH1000 (*41*) was grown to Mid-Log phase in brain heart infusion medium (BHI, Roth, Karlsruhe, Germany) at 37°C with shaking (120 rpm), collected by centrifugation, washed with sterile PBS, and diluted to the required concentration. The number of viable bacteria was determined after serial diluting and plating on BHI-agar.

#### Mice and infection

A previously described chronic renal abscess infection model (*5*) has been used in this study. It is known that *S. aureus* strain SH1000 produces renal abscesses after intravenous inoculation (*5*). Pathogen-free, 10 weeks-old C57BL/6 female mice were purchased from Harlan-Winkelmann (Envigo, Netherlands). All animals were provided with food and water ad libitum, and housed in groups of up to 5 mice per cage in individually ventilated cages. Statistical power analysis with anticipated incidence of 10% and 90% for control and treated mice, respectively, indicated a sample size of five mice per group. Mice were infected with 5×10^7^ CFU of *S. aureus* in 100 μl of PBS via a tail vein and monitored on a daily basis for weight loss and sign of pain or distress. At specified times of infection, mice were sacrificed by CO_2_ asphyxiation and the bacterial load was enumerated in kidney homogenates by plating 10-fold serial dilutions on blood agar plates. Spleens were removed, transformed in a single cell suspension and further processed for FACS and proliferation assays.

In vaccination experiments, infected mice were injected intraperitoneally at day 14 of infection with 10^8^ heat-killed bacteria of *S. aureus* strain SH1000 or *S. pyogenes* strain A20 in 200 μl of PBS that were prepared by heating a bacterial suspension at 60°C for 1 h. At 12 h postchallenge, mice were sacrificed and peritoneal exudate cells (PEC) were isolated from infected mice by lavage of the peritoneal cavity with 2 ml sterile PBS. The lavage fluid was centrifuged, supernatants stored at −20°C for subsequent cytokine analysis, and PEC resuspended in complete RPMI, stained and analyzed by flow cytometry (see below).

Animal experiments were performed in strict accordance with the German regulations of the Society for Laboratory Animal Science (GV-SOLAS) and the European Health Law of the Federation of Laboratory Animal Science Associations (FELASA). All experiments were approved by the ethical board Niedersächsisches Landesamt für Verbraucherschutz und Lebensmittelsicherheit, Oldenburg, Germany (LAVES; permit N. 18/2798).

#### Flow cytometry analysis

Cells were incubated with purified rat anti-mouse CD16/CD32 (BD Biosciences) for 5 min to block Fc receptors and then stained with antibodies against CD11b (BioLegend), Ly6C (BioLegend), Ly6G (Miltenyi Biotec), F4/80 (BD Biosciences) for 20 min at 4°C. Labeled cells were measured by flow cytometry using a BDTM LSR II flow cytometer (BD Biosciences) and analyzed by FlowJo software.

#### Proliferation assay

Spleen cells were seeded into 96-well flat-bottom plates at 5 ×10^5^ cells/well in 100 μl of complete RPMI medium and stimulated with 2 μg/ml of anti-CD3/anti-CD28 antibodies (Sigma-Aldrich) at 37°C and 5% CO_2_ . After 3 days of incubation, the cells were pulsed with 1 μCi ^3^H-thymidine (Amersham) and harvested 18 h later on Filtermats A (Wallac) using a cell harvester (Inotech). The amount of 3H-thymidine incorporation was measured in a gamma scintillation counter (Wallac 1450; MicroTrilux).

#### Statistical analyses

All data were analyzed with GraphPad Prism Software. Comparisons between several groups were made using a parametric ANOVA test with Tukey’s multiple comparisons test. Comparison between two groups was performed using a Student’s t-test. P values <0.05 were considered significant.

## Supplementary Materials

### Supplementary Methods

#### Mathematical model

The mathematical model was implemented and simulated in MATLAB, see www.mathworks.com.

##### Fitting curves and standard deviation of parameters

The unknown parameters in the model were estimated in three steps. First, the carrying capacity of bacteria κ was estimated based on our previously reported experimental results from B and T cells-deficient RAG2^−/−^ mice (only innate immunity present), which were initially infected with 7 × 10^7^ CFU of *S. aureus* strain SH1000 (same strain as used in the study here) and showed a nearly constant level of *S. aureus* in kidneys from day 7 till day 56 (*5*). The mean of these data points was taken as the bacterial carrying capacity. The carrying capacity of T cells (parameter *K_T_*) was assumed to be of the same order of magnitude as the bacterial carrying capacity. Secondly, the growth rate of bacteria was estimated by solving the logistic growth equation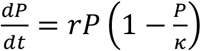 for r

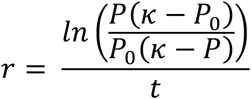

where κ is the bacterial carrying capacity, *P*_0_ is the initial inoculation number of bacteria and P is the bacterial CFU at time t. Our previous experimental data showed that the bacterial loads in B and T cells-deficient RAG2^−/−^ mice reached 75% of the carrying capacity on day 2 and fluctuated afterwards, after intravenous inoculation with 7×10^7^ CFU of *S. aureus*. Assuming that the bacterial load reached 75% of the carrying capacity by day 1 or day 2, we determined the high and low boundary of *r* to be 0.636 and 0.318, respectively. The average of the low and high boundary was taken as the bacterial growth rate *r_b_* of the model. Finally, our previously reported data of bacterial loads and absolute numbers of T cells in time from immunocompetent mice with chronic *S. aureus* SH1000-infection and renal abscesses (*5*) were used to fit the rest of unknown parameters. The fitting process used a Markov Chain Monte Carlo version of Differential Evolution algorithm *(42)*. Parameter *c_T_*, representing T-cell suppression locally (at the site of infection which is the kidneys for the chronic infection here) appeared much smaller than other parameters, in particular parameter Θ (representing T-cell suppression by MDSCs systemically, namely distant from the site of infection, such as in spleen). We tested, in a second study, the possibility of fitting the same data with *c_T_* = 0. The fit quality remained the same; therefore we concluded that *c_T_* is zero and that the main T-cell suppression by MDSCs is exerted systemically. This deduction accords with our previous experimental reports which showed major T-cell suppression by MDSCs in spleens of mice that had chronic *S. aureus* infection in kidneys (*6*, *13*). Fitted parameter values are shown in table S1.

##### Scaling the model

*For the analytical solutions and analytical stability analysis the calculations were made feasible by using an analytically amenable ODE model, which approximates the original ODE as described in Eq. (1)-(2).*

With the following change of variables we non-dimensionalize the ODE model (1)-(2):

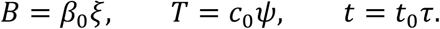

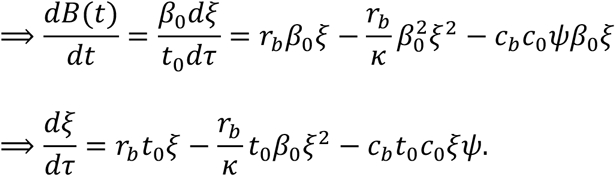

We impose that the coefficients of *ξ*, *ξ*^2^and *ξ*ψ are equal 1. Then

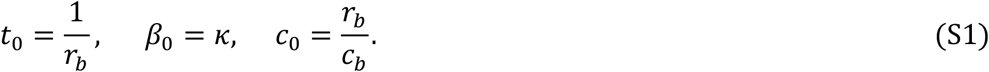

Therefore the new, non-dimensionalized equation for bacteria is

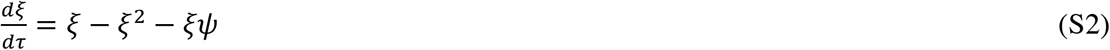

For the equation *T*(*t*) we have the non-dimensionalized calculations:

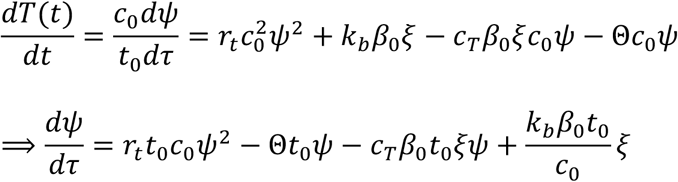

Substitution of the *t*_0_, β_0_ and *c*_0_ (found in Eq. (S1)) gives the scaled equation for T:

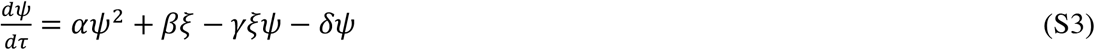

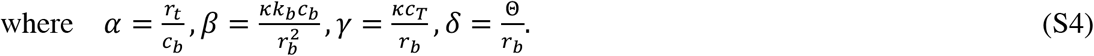

##### Calculation of equilibrium points

From Eq. (S2) we have:

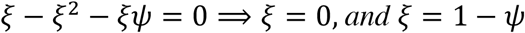

- For *ξ* = 0 in Eq. (S3) we have:

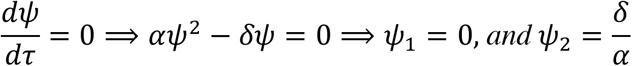
- For *ξ* = 1 − *ψ* in Eq. (S3) in steady state we conclude that:

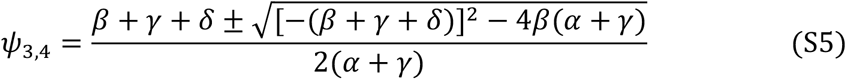

Consequently the system has four equilibrium points in total:

- (*ξ*_2_, *ψ*_2_) = (0,0)
- 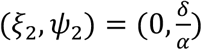
- (*ξ*_3,4_, *ψ*_3,4_) = (1 − *ψ*_3,4_, *ψ*_3,4_) where *ψ*_3,4_ is as shown above.

##### Existence of *ψ*_3,4_

Equilibrium points *ψ*_3,4_ exist only when *Δ* = (*β* + *γ* + *δ*)^2^ − 4*β*(*α* + *γ*) ≥ 0, i.e. only when *ψ*_3,4_ have no imaginary part. Equivalently

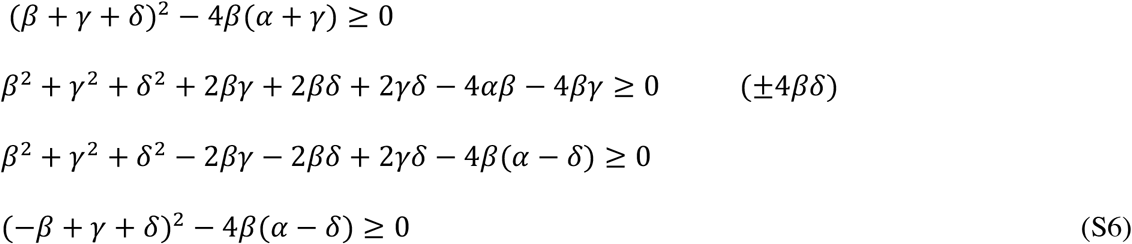

According to the sign of the term (*α* − *δ*) in condition (S6), we investigate when the equilibrium points *ψ*_3,4_ exist.

- If *α* − *δ* ≤ 0 ⟹ *δ* ≥ *a* then the equilibria *ψ*_3,4_ exist.
- If 0 < *δ* < *α* then we reformulate *Δ* as follows:

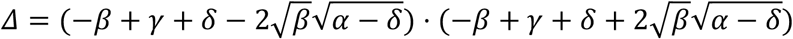
  a. If −*β* + *γ* + *δ* ≥ 0 then for existence of *ψ*_3,4_ we require 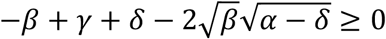. Then:

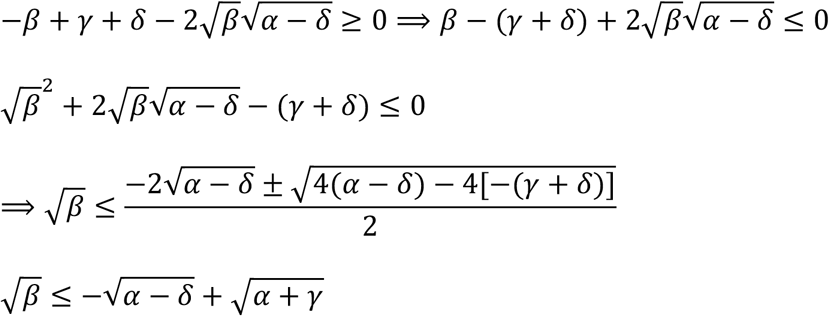 **Note:** The solution 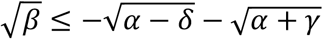 is exempt because by definition 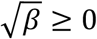.
  b. If −*β* + *γ* + *δ* ≤ 0 then for existence of *ψ*_3,4_ we require 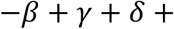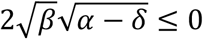. Then:

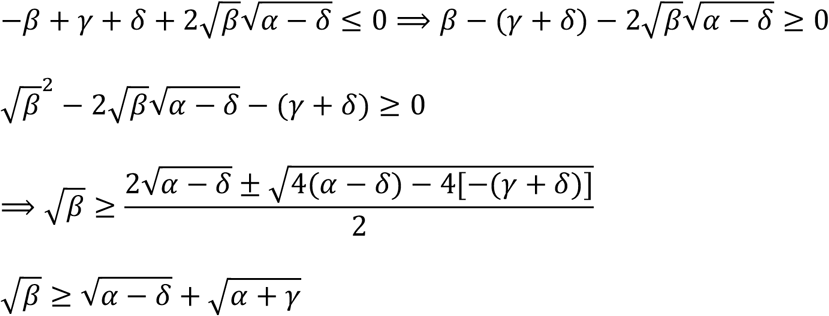 **Note:** The solution 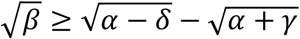 is trivially exempt.

##### Local stability analysis of equilibrium points

Jacobian Matrix 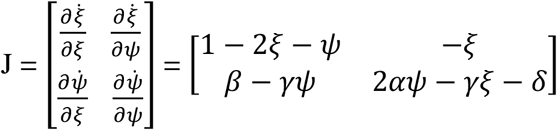

Evaluation of Jacobian matrix in the trivial equilibrium point (*ξ*_2_, *ψ*_2_):

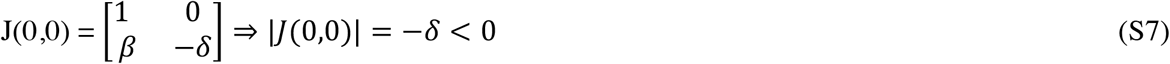

From linear algebra, it is known that if J is a 2×2 matrix and λ_1_, λ_2_ its eigenvalues, then the determinant of J, |J|= *λ*1 × *λ*2. Since the determinant of the Jacobian matrix in Eq. (S7) is negative (i.e. det = −δ <0), it means that the eigenvalues are of a different sign, and hence the trivial equilibrium point is unstable (i.e. saddle point, which is always unstable).

For the equilibrium point 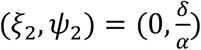 we have: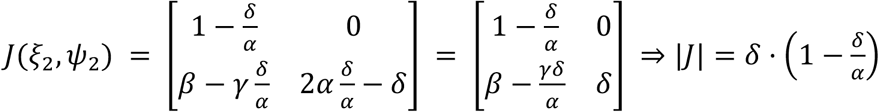 for the equilibrium point (*ξ*_2_, *ψ*_2_). The sign of the determinant depends on the term 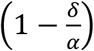. When *δ* > *α*, then |*J*| < 0 (hence one positive and one negative eigenvalue), meaning that the equilibrium point is saddle. This becomes unstable when *δ* < *α*. Next, we evaluate the Jacobian matrix in the equilibrium points (*ξ*_3,4_, *ψ*_3,4_), by using *ξ* = 1 − *ψ* ≥ 0,

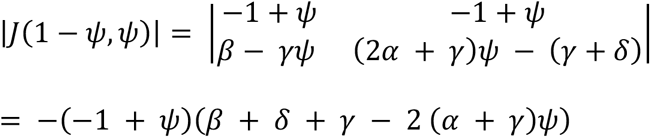

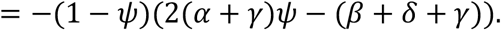

We obtain the trace of the Jacobian matrix, tr(J), to determine the type of stability of equilibrium points (*ξ*_3,4_, *ψ*_3,4_),

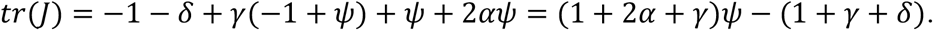

We first need to find some critical values for ψ:

- If |*J*(1 − *ψ*, *ψ*)| = 0, then

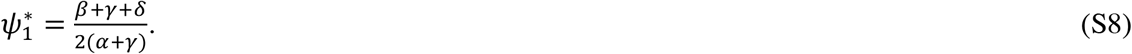

We know that the term (1 − *ψ*) equals ξ, which represents the bacteria, and hence *ξ* = 1 − *ψ* ≥ 0.

- If *tr*(*J*) = 0, then

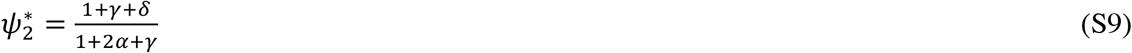

From the critical points found in Eqs. (S8) and (S9), the stability of the equilibrium points (*ξ*_3,4_, *ψ*_3,4_) = (1 − *ψ*_3,4_, *ψ*_3,4_) can be classified as follows:

***Stable node or spiral (Represents the chronic phase):***

- 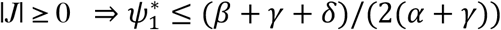
- 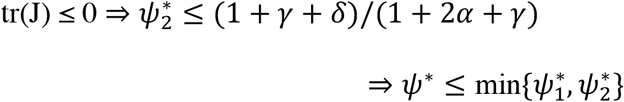

***Saddle equilibrium point:***

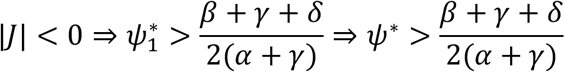

***Unstable node or spiral:***

- 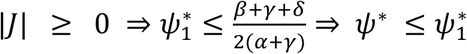
- 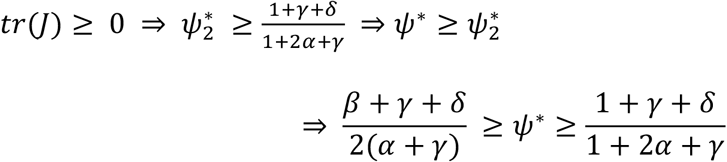

However, since from fitting results (table S1) *c_T_* = 0, we conclude that

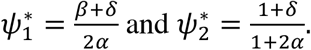

Assuming 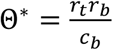 and substituting α, β, and δ from Eqs. (S4),

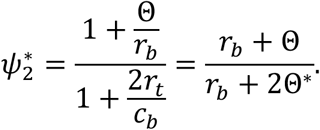

Since ξ and ψ are normalized, ξ, ψ ≥ 0 and therefore 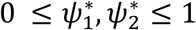. Hence 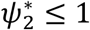 resulting in

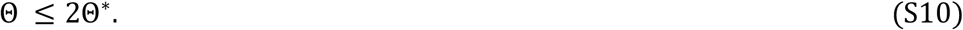

Now, 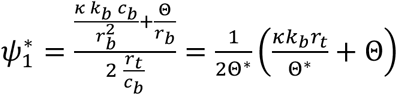.

From Eq. (S10)

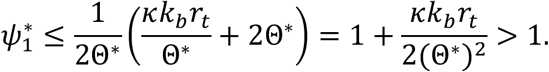

As a consequence, 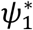 is rejected and the stability analysis for the equilibrium points (*ξ*_3,4_, *ψ*_3,4_) = (1 − *ψ*_3,4_, *ψ*_3,4_) can be updated as follows:

- ***Stable node or spiral (Represents the chronic phase):***

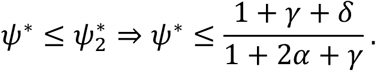
- ***Unstable node or spiral:***

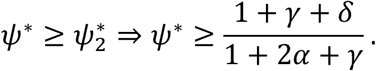

## Supplementary figures

**Fig. S1.**
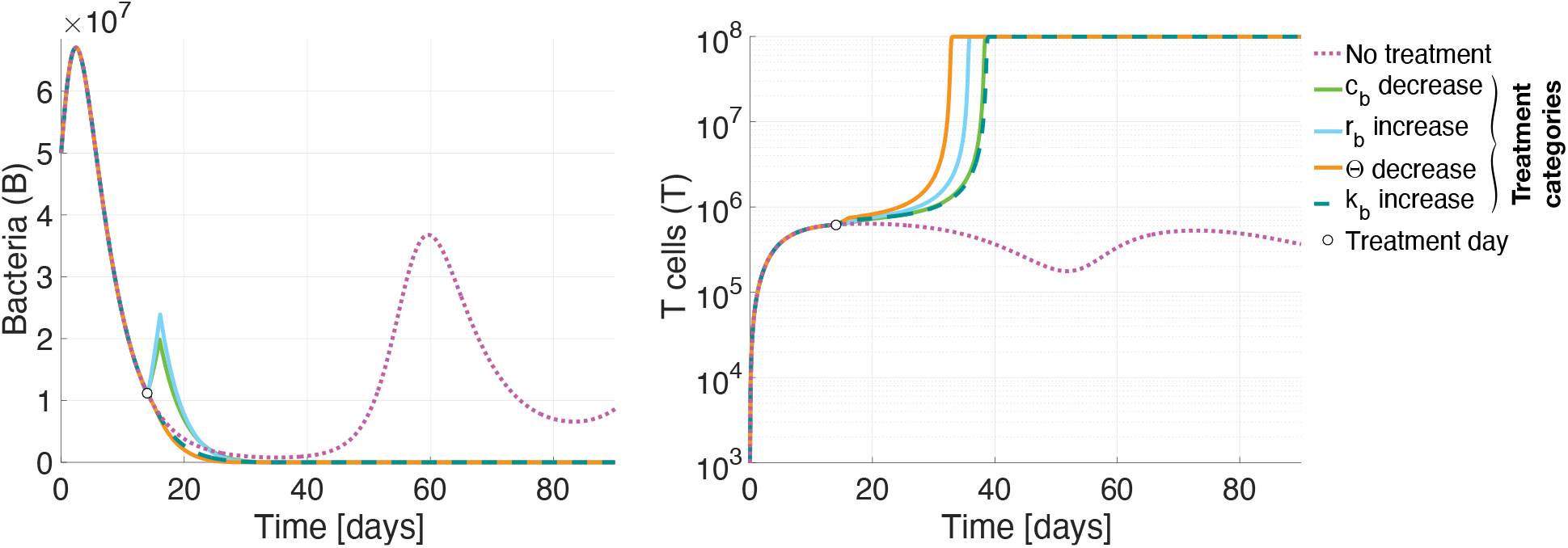
Perturbation strategies suggest eradication of *S. aureus in silico*. Progression of *S. aureus* infection without any perturbation treatment results in a mathematically-characterized stable state and in a clinically-characterized persistent (chronic) *S. aureus* infection. When the chronic infection system is perturbed with treatments that (i) decrease host aggression against bacteria (*c_b_*), for example via immunosuppressant drugs, (ii) increase bacterial growth (*r_b_*), (iii) reduce MDSC-mediated immunosuppression (Θ), for instance via gemcitabine chemotherapeutic drug, or (iv) boosting adaptive immunity of the host (*k_b_*), such as via administration of heat-killed bacterial antigens, *S. aureus* is eradicated and the host is cured. The plotted bacterial and T-cell dynamics in time are the numerical solutions of the ODE system (Eqs. (1)-(2)). Treatments were applied *in silico* at day 14 of infection (white bullet), namely when infection was already chronic, by decreasing or increasing the fitted value of the parameter of interest (table S1) for a perturbation window of 12 hours up to two days.

**Fig. S2.**
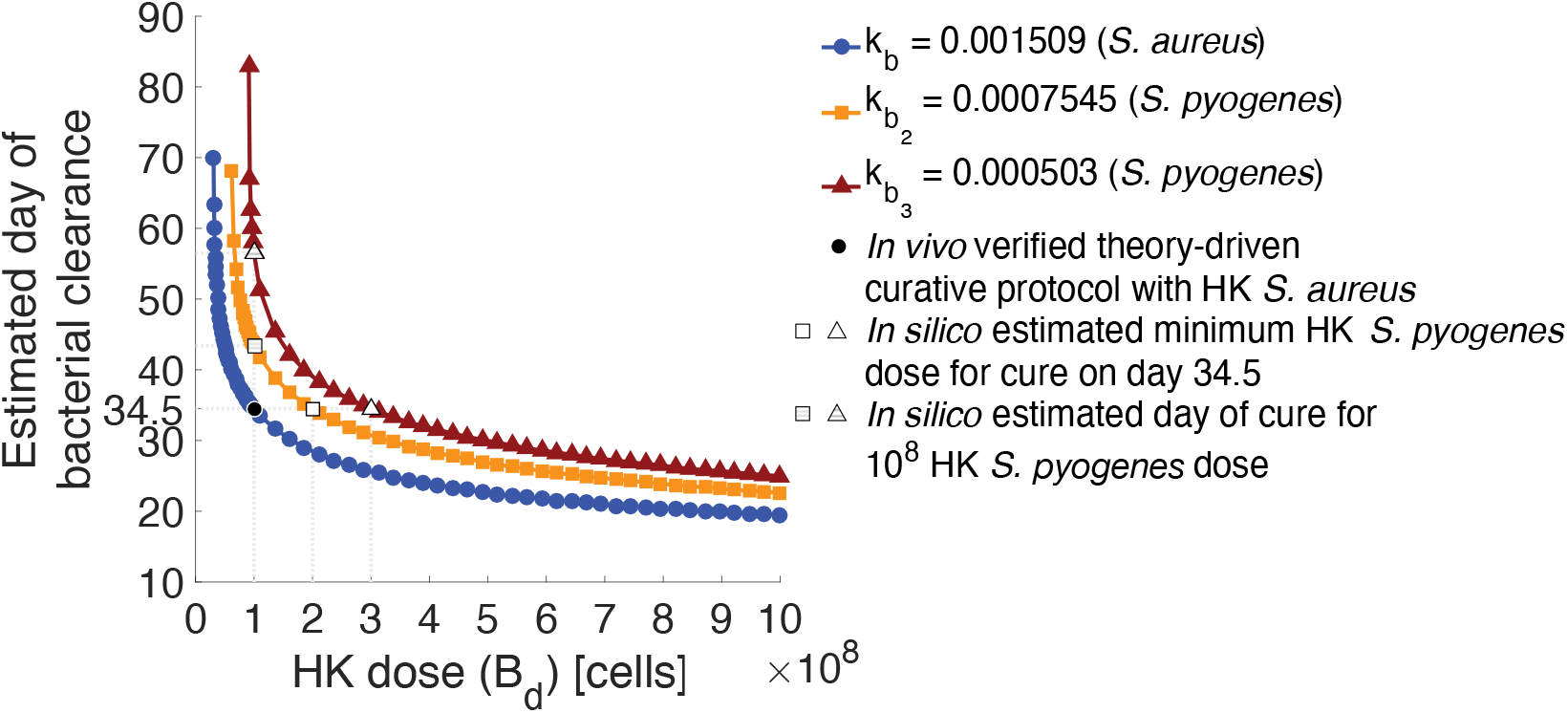
Time of bacterial clearance for varying HK-doses and three values of T-cell stimulatory parameter *k_b_ in silico*. Stimulation of T cells by HK *S. aureus* cells at day 14 happens with rate *k_b_* = 0.001509 as defined during the fitting process (table S1). Since primary exposure to *S. aureus* already occurs upon inoculation at day 0, HK *Streptococcus pyogenes* at day 14 of infection stimulates the immune system at a weaker and slower rate compared to HK *S. aureus*, assumed as *k_b2_* = *k_b_*/2 = 0.0007545 or *k_b3_* = *k_b_*/3 = 0.000503. Simulation of HK treatments was done by adding the term *k_b_B_d_* to the T-cell ODE at day 14 of infection for a perturbation window of half day (*Materials and Methods*), where *k_b_* the immunostimulatory parameter varying for three different values (*k_b_*, *k_b_*/2, *k_b_*/3) and *B_d_* the HK-doses varying in the range [4×10^7^,10^9^]. The estimated day of clearance was defined as the first time-point when bacterial numbers <0.000001.

## Supplementary table

**Table S1.** Model parameter values as used for model analysis.

| Parameter | Description | Value, [Confidence Interval]* | Unit |
| --- | --- | --- | --- |
| $r_b$ | Bacterial growth rate (in the presence of innate immunity) | 0.477 | days<sup>-1</sup> |
| $\kappa$ | Carrying capacity of bacteria | $1.132 \times 10^8$ | cells |
| $c_b$ | Rate of bacterial containment by T cells (per cell) | $9.937 \times 10^{-7}$, [$8.65 \times 10^{-7}$, $1.25 \times 10^{-6}$] | days<sup>-1</sup> |
| $r_t$ | T-cell proliferation rate (per cell) | $2.0955 \times 10^{-7}$, [$1.04 \times 10^{-7}$, $2.94 \times 10^{-7}$] | days<sup>-1</sup> |
| $k_b$ | T-cell activation and recruitment rate | 0.001509, [0.0011, 0.0017] | days<sup>-1</sup> |
| $c_T$ | Local T-cell suppression rate (per cell) | 0 | days<sup>-1</sup> |
| $\Theta$ | Systemic MDSC-mediated suppression rate on T cells | 0.14393, [0.072, 0.18] | days<sup>-1</sup> |
| $K_T$ | Carrying capacity of T cells | $10^8$ | cells |

\* In square brackets is the 95% confidence interval of the parameters as derived by a Markov Chain Monte Carlo version of Differential Evolution algorithm (42).

## Acknowledgments

The authors thank Sabine Lehne and Lothar Gröbe for excellent technical assistance, and Drs. Anna Ntalli, Heiko Enderling, Andreas Papaxenopoulos, Yasmina Marin-Felix, Andreas I. Andreou and Kamila Tomoko Yuyama for useful comments on a draft of this article.

## Author contributions

LP, HH, SK, GZ, and MMH designed the study, developed the methodology and interpreted the results. GZ performed the fitting process. LP performed the model implementation and simulations, and analyzed the results. LP and EM designed and conducted the experiments. LP, KKD and IS analyzed the experimental results. HH and MMH supervised the study. All authors wrote the paper and approved the final version of the manuscript.

## Data and materials availability

Original experimental data as well as MATLAB codes are provided in the manuscript or in the Supplementary Materials.

## REFERENCES

1. L. Thomer, O. Schneewind, D. Missiakas, Pathogenesis of *Staphylococcus aureus* bloodstream infections. Annu. Rev. Pathol. Mech. Dis. 11, 343–364 (2016).

2. S. D. Kobayashi, N. Malachowa, F. R. DeLeo, Pathogenesis of *Staphylococcus aureus* abscesses. Am. J. Pathol. 185, 1518–1527 (2015).

3. B. L. Hahn, P. G. Sohnle, Direct translocation of staphylococci from the skin surface to deep organs. Microb. Pathog. 63, 24–29 (2013).

4. S. V. Kazakova, J. C. Hageman, M. Matava, A. Srinivasan, L. Phelan, B. Garfinkel, T. Boo, S. McAllister, J. Anderson, B. Jensen, D. Dodson, D. Lonsway, L. K. McDougal, M. Arduino, V. J. Fraser, G. Killgore, F. C. Tenover, S. Cody, A clone of methicillin resistant *Staphylococcus aureus* among professional football players. N. Engl. J. Med. 352, 468–475 (2005).

5. C. Ziegler, O. Goldmann, E. Hobeika, R. Geffers, G. Peters, E. Medina, The dynamics of T cells during persistent *Staphylococcus aureus* infection: from antigen-reactivity to in vivo anergy. EMBO Mol. Med. 3, 652–666 (2011).

6. C. Tebartz, S. A. Horst, T. Sparwasser, J. Huehn, A. Beineke, G. Peters, E. Medina, A major role for myeloid-derived suppressor cells and a minor role for regulatory T cells in immunosuppression during *Staphylococcus aureus* infection. J. Immunol. 194, 1100–1111 (2015).

7. M. Ost, A. Singh, A. Peschel, R. Mehling, N. Rieber, D. Hartl, Myeloid-derived suppressor cells in bacterial infections. Front. Cell. Infect. Microbiol. 6(2016).

8. E. Medina, D. Hartl, Myeloid-derived suppressor cells in infection: a general overview. J. Innate Immun. 10, 407–413 (2018).

9. C. Agrati, A. Sacchi, V. Bordoni, E. Cimini, S. Notari, G. Grassi, R. Casetti, E. Tartaglia, E. Lalle, D’ Abramo, C. Castilletti, L. Marchioni, Y. Shi, A. Mariano, J.-W. Song, J.-Y. Zhang, F.-S. Wang, C. Zhang, G. M. Fimia, M. R. Capobianchi, M. Piacentini, A. Antinori, E. Nicastri, M. Maeurer, A. Zumla, Expansion of myeloid-derived suppressor cells in patients with severe coronavirus disease (COVID-19). Cell Death Differ. 27, 3196–3207 (2020).

10. E.-K. Vetsika, A. Koukos, A. Kotsakis, Myeloid-derived suppressor cells: major figures that shape the immunosuppressive and angiogenic network in cancer. Cells 8, 1647 (2019).

11. C. Guo, F. Hu, H. Yi, Z. Feng, C. Li, L. Shi, Y. Li, H. Liu, X. Yu, H. Wang, J. Li, Z. Li, X.-Y. Wang, Myeloid-derived suppressor cells have a proinflammatory role in the pathogenesis of autoimmune arthritis. Ann. Rheum. Dis. 75, 278–285 (2016).

12. O. Goldmann, E. Medina, *Staphylococcus aureus* strategies to evade the host acquired immune response. Int. J. Med. Microbiol. 308, 625–630 (2018).

13. O. Goldmann, A. Beineke, E. Medina, Identification of a novel subset of myeloid-derived suppressor cells during chronic staphylococcal infection that resembles immature eosinophils. J. Infect. Dis. 216, 1444–1451 (2017).

14. G. A. Noskin, R. J. Rubin, J. J. Schentag, J. Kluytmans, E. C. Hedblom, M. Smulders, E. Lapetina, E. Gemmen, The burden of *Staphylococcus aureus* infections on hospitals in the United States: an analysis of the 2000 and 2001 Nationwide Inpatient Sample Database. Arch. Intern. Med. 165, 1756–1761 (2005).

15. E. Y. Klein, W. Jiang, N. Mojica, K. K. Tseng, R. McNeill, S. E. Cosgrove, T. M. Perl, National costs associated with methicillin-susceptible and methicillin-resistant *Staphylococcus aureus* hospitalizations in the United States, 2010–2014. Clin. Infect. Dis. 68, 22–28 (2018).

16. F. Cedeira, F. Cedeira, E. M. Pagano, G. Castañeda, R. Rivadera, Bilateral nephrectomy as an extreme measure management for methicillin-resistant *Staphylococcus aureus* (MRSA): a case report. Urol. Case Rep. 33, 101373 (2020).

17. C. Hogea, T. Van Effelterre, C. J. Acosta, A basic dynamic transmission model of *Staphylococcus aureus* in the US population. Epidemiol. Infect. 142, 468–478 (2014).

18. E. McBryde, A. Pettitt, D.L.S. McElwain, A stochastic mathematical model of methicillin resistant *Staphylococcus aureus* transmission in an intensive care unit: predicting the impact of interventions. J. Theor. Biol. 245, 470–481 (2007).

19. W.-S. Choi, N. Son, J.-I. Cho, I.-S. Joo, J.-A. Han, H.-S. Kwak, J.-H. Hong, S. H. Suh, Predictive model of *Staphylococcus aureus* growth on egg products. Food Sci. Biotechnol. 28, 913–922 (2019).

20. A. Best, J. Jubrail, M. Boots, D. Dockrell, H. Marriott, A mathematical model shows macrophages delay *Staphylococcus aureus* replication, but limitations in microbicidal capacity restrict bacterial clearance. J. Theor. Biol. 497, 110256 (2020).

21. S. K. Shukla, T. C. Carter, Z. Ye, M. Pantrangi, W. E. Rose, Modeling of effective antimicrobials to reduce *Staphylococcus aureus* virulence gene expression using a two-compartment hollow fiber infection model. Toxins 12, 69 (2020).

22. M. Schmaler, N. J. Jann, F. Ferracin, R. Landmann, T and B cells are not required for clearing *Staphylococcus aureus* in systemic infection despite a strong TLR2–MyD88-dependent T cell activation. J. Immunol. 186, 443–452 (2011).

23. H. K. Kim, V. Thammavongsa, O. Schneewind, D. Missiakas, Recurrent infections and immune evasion strategies of *Staphylococcus aureus*. Curr. Opin. Microbiol. 15, 92–99 (2012).

24. S. Khailaie, F. Bahrami, M. Janahmadi, P. Milanez-Almeida, J. Huehn, M. Meyer-Hermann, A mathematical model of immune activation with a unified self-nonself concept. Front. Immunol. 4, 474 (2013).

25. C. H. Kim, Homeostatic and pathogenic extramedullary hematopoiesis. J. Blood Med. 1, 13 (2010).

26. C. E. Heim, D. Vidlak, T. D. Scherr, J. A. Kozel, M. Holzapfel, D. E. Muirhead, T. Kielian, Myeloid-derived suppressor cells contribute to *Staphylococcus aureus* orthopedic biofilm infection. J. Immunol. 192, 3778–3792 (2014).

27. C. Ma, T. Kapanadze, J. Gamrekelashvili, M. P. Manns, F. Korangy, T. F. Greten, Anti-Gr-1 antibody depletion fails to eliminate hepatic myeloid-derived suppressor cells in tumor-bearing mice. J. Leukoc. Biol. 92, 1199–1206 (2012).

28. S. Ostrand-Rosenberg, P. Sinha, Myeloid-derived suppressor cells: linking inflammation and cancer. J. Immunol. 182, 4499–4506 (2009).

29. J. S. Ayres, Surviving COVID-19: a disease tolerance perspective. Sci. Adv. 6, eabc1518 (2020).

30. K. A. Anderson, J. W. McAninch, Renal abscesses: classification and review of 40 cases. Urology 16, 333–338 (1980).

31. T. J. Yoon, J. Y. Kim, H. Kim, C. Hong, H. Lee, C.-K. Lee, K. H. Lee, S. Hong, S.-H. Park, Anti-tumor immunostimulatory effect of heat-killed tumor cells. Exp. Mol. Med. 40, 130–144 (2008).

32. A. d. A. Cassado, M. R. D’Império Lima, K. R. Bortoluci, Revisiting mouse peritoneal macrophages: heterogeneity, development, and function. Front. Immunol. 6, 225 (2015).

33. E. E. B. Ghosn, A. A. Cassado, G. R. Govoni, T. Fukuhara, Y. Yang, D. M. Monack, K. R. Bortoluci, S. R. Almeida, L. A. Herzenberg, L. A. Herzenberg, Two physically, functionally, and developmentally distinct peritoneal macrophage subsets. Proc. Natl. Acad. Sci. U.S.A 107, 2568–2573 (2010).

34. E. Takenaka, A. Van Vo, Y. Yamashita-Kanemaru, A. Shibuya, K. Shibuya, Selective DNAM-1 expression on small peritoneal macrophages contributes to CD4+ T cell costimulation. Sci. Rep. 8, 1–8 (2018).

35. S. Holtfreter, J. Jursa-Kulesza, H. Masiuk, N. Verkaik, C. de Vogel, J. Kolata, M. Nowosiad, L. Steil, W. van Wamel, A. van Belkum, U. Völker, S. Giedrys-Kalemba, B. M. Broker, Antibody responses in furunculosis patients vaccinated with autologous formalin-killed *Staphylococcus aureus*. Eur. J. Clin. Microbiol. Infect. Dis. 30, 707 (2011).

36. D. Missiakas, O. Schneewind, *Staphylococcus aureus* vaccines: deviating from the carol. J. Exp. Med. 213, 1645–1653 (2016).

37. V. Vyas, T. Endy, A Rare case of prostatic and bilateral renal abscesses caused by community-acquired methicillin-resistant *Staphylococcus aureus* infection. Cureus 12(2020).

38. T. Fiedler, T. Köller, B. Kreikemeyer, *Streptococcus pyogenes* biofilms—formation, biology, and clinical relevance. Front. Cell. Infect. Microbiol. 5, 15 (2015).

39. J. Stanford, C. Stanford, J. Grange, Immunotherapy with *Mycobacterium vaccae* in the treatment of tuberculosis. Front Biosci. 9, 1701–1719 (2004).

40. M. I. Gröschel, S. A Prabowo, P.-J. Cardona, J. L. Stanford, T. S. Van der Werf. Therapeutic vaccines for tuberculosis—a systematic review. Vaccine. 32, 3162–3168 (2014).

41. M. Jonsson, S. Arvidson, S. Foster, A. Tarkowski, Sigma factor B and RsbU are required for virulence in *Staphylococcus aureus*-induced arthritis and sepsis. Infect. Immun. 72, 6106–6111 (2004).

42. C. J. Ter Braak, A markov chain monte carlo version of the genetic algorithm differential evolution: easy bayesian computing for real parameter spaces. Stat. Comput. 16, 239–249 (2006).

